# Metabolic control of enteroendocrine cell fate through a redox state sensor CtBP

**DOI:** 10.1101/2025.06.30.662346

**Authors:** Bohdana M. Rovenko, Kari Moisio, Cornelia Biehler, Mykhailo Girych, Atte Hallasaari, Onur Deniz, Sarah Bluhm, Arto Viitanen, Krista Kokki, Gaia Fabris, Yi Yang, Valentin Cracan, Irene Miguel-Aliaga, Ville Hietakangas

## Abstract

Enteroendocrine (EE) cells monitor the intestinal nutrient composition and consequently control organismal physiology through hormonal signaling. In addition to the immediate effects on hormone secretion, nutrients influence EE cell abundance by affecting the determination and maintenance of cell fate. EE cells are known to import and respond to dietary sugars, but how the sugar-induced changes in the intracellular metabolic state are sensed to control the immediate and long-term responses of EE cells, remains poorly understood. We report that the NADH binding transcriptional cofactor C-terminal binding protein (CtBP) acts at the interface between nutrient sensing and fate regulation of *Drosophila* larval EE cells, thus controlling organismal energy metabolism and survival on a high sugar diet. CtBP dimerization in EE cells is regulated through the redox balance of nicotinamide cofactors controlled by glycolysis and pentose phosphate pathway, allowing EE cells sense their internal metabolic state in response to sugar catabolism. CtBP interacts with the EE cell fate determining transcription factor Prospero through a conserved binding motif and binds to genomic targets controlling EE cell fate and size, such as components of Notch and insulin/mTOR pathways. Collectively, our findings uncover a modality where changes in intracellular redox state serve as an instructive signal to control EE cell function to globally control organismal homeostasis.

**GRAPHICAL ABSTRACT:** 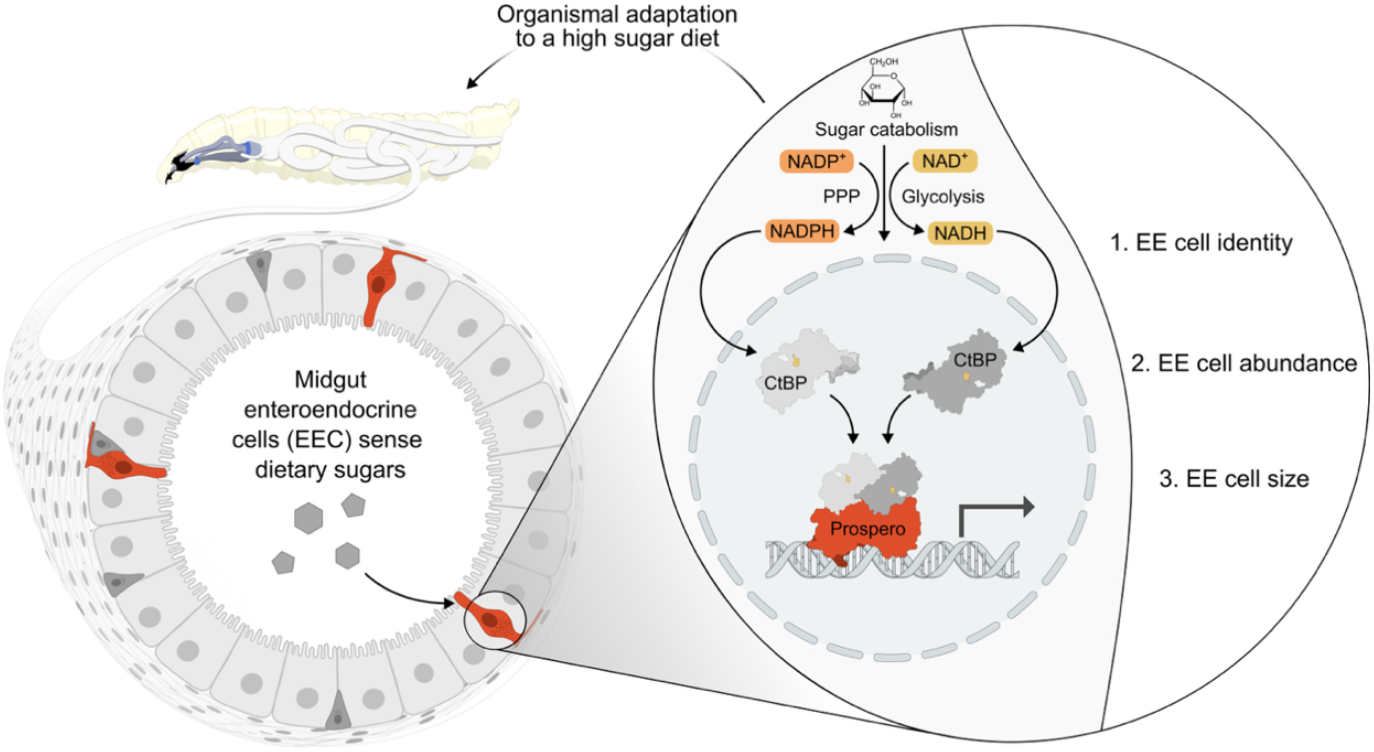

## INTRODUCTION

Physiological functions, such as metabolic homeostasis, are closely controlled in response to nutrient intake. Sensing of nutrients starts at the first point of contact with the ingested food. Enteroendocrine (EE) cells are key mediators of this early nutrient sensing, as they are distributed throughout the gastrointestinal epithelium to monitor intestinal nutrient content. EE cells are comprised of different subtypes, with distinct peptide hormones and nutrient responsiveness and their physiological functions are well conserved in metazoan species^1-3^. EE cells have well-established roles in glucose homeostasis. For example, glucagon-like peptide-1 (GLP-1) is secreted by so-called L-cells abundant in mammalian ileum and colon and it controls systemic glucose homeostasis by stimulating insulin secretion in pancreatic β-cells^4^. In *Drosophila*, similar incretin effects are mediated by neuropeptide F (NPF), which is secreted by midgut EE-cells in a sugar-responsive manner and promotes systemic insulin signaling through its receptor in the insulin-producing cells^5^. Despite these insights, questions regarding the molecular mechanisms of EE cell nutrient sensing and roles of EE hormones controlling metabolism in response to specific macronutrients remain insufficiently understood.

To be able to monitor changes in nutrient availability, EE cells express a variety of membrane proteins, including nutrient transporters and G protein-coupled receptors, which regulate hormone release^2^. For example, the sugar-responsive secretion of *Drosophila* NPF requires expression of Sugar transporter1 (Sut1) in EE cells^5^, while the secretion of another EE hormone, Bursicon α, is controlled by Glucose transporter 1 (Glut1)^6^. How the imported sugars and their downstream metabolites are sensed intracellularly and how they impact EE cell functions remain, however, poorly understood. In addition to the immediate effects on hormone secretion, nutrients can have long-term impact on EE cells through fate regulation. For example, dietary cholesterol increases EE cell numbers in adult *Drosophila* by influencing Notch signaling, which controls EE cell differentiation^7^. Notably, in obese human patients the density of GLP-1 secreting L-cells is higher in individuals with a high fat diet, suggesting that dietary lipids may have conserved role in controlling EE cell abundance^8^. The molecular mechanisms mediating long-term adaptive nutrient responses on EE cells remain mostly unresolved.

Redox balance, the relative state of reduction and oxidation of metabolites, is a critical determinant of cellular homeostasis^9^. Among the various redox-active compounds in the cell, nicotinamide cofactors constitute a buffering capacity for redox potential, enabling dynamic adaptation of cellular metabolism in response to changes in nutrient availability^10,11^. Upon glucose catabolism, nicotinamide dinucleotide NAD^+^ is reduced through the reactions of glycolysis and the tricarboxylic acid (TCA) cycle, while its phosphorylated counterpart NADP^+^ stores reductive potential released by the pentose phosphate pathway. Beyond enabling reactions requiring oxidative and reductive transformations, these cofactors serve as signaling molecules by binding to and modulating the activity of regulatory proteins^9,12^. A well-characterized example of such proteins is the Sirtuin family of NAD^+^-dependent deacetylases, which regulate physiological processes in energy metabolism, stress resistance, and aging^13,14^. Significantly less attention has been given to the physiological roles of C-terminal binding protein (CtBP), a transcriptional corepressor whose dimerization and transcriptional activity is controlled by binding of nicotinamide cofactors^12,15,16^. *In vitro* studies have shown that CtBP binds to the reduced form of nicotinamide dinucleotide (NADH) with >100-fold higher affinity^15^ than to the oxidized form NAD^+^. This suggests that CtBP activity might be coupled to the cellular redox balance, and consequently the metabolic state of the cell, albeit opposing arguments have also been presented due to higher relative abundance of NAD^+17^. While mouse CtBP paralog CtBP2 is known to repress Forkhead box O1 (FoxO1)-mediated hepatic gluconeogenesis^18^, other physiological roles of CtBP in organismal nutrient sensing and metabolic regulation remain poorly characterized.

Here we identify a regulatory axis that couples molecular sensing of EE cell intrinsic sugar catabolism with nutrient-responsive control of EE cell abundance. We demonstrate that the conserved transcriptional corepressor C-terminal binding protein (CtBP) has a necessary role in regulating EE cells in *Drosophila* larvae, contributing to the systemic control of metabolism and maintaining viability of animals exposed to a high sugar diet. Nutrient-induced changes in EE cell redox balance of nicotinamide cofactors NAD(H) and NADP(H) control CtBP oligomerization status, thus allowing EE cells to sense their intrinsic metabolic state by this molecular switch. CtBP associates with target genes controlling EE cell size and fate, including components of the insulin/mTOR and Notch signaling pathways. CtBP interacts with the EE cell fate regulator Prospero in the EE cells and it shares a significant portion of direct targets with Pros, including an EE cell fate regulator *split ends (spen)*. Our findings are consistent with a model that CtBP serves as a metabolic state-dependent modulator of EE cell fate, allowing dynamic control of EE cell abundance in response to nutrient intake.

## RESULTS

### CtBP regulates Drosophila energy metabolism and is necessary for sugar tolerance

The metabolic pathways influencing redox balance of nicotinamide cofactors have an essential role in mediating the metabolic adaptation to a high sugar diet in *Drosophila* larvae^19,20^. Therefore, we wanted to better understand the possible role of nicotinamide cofactor-binding proteins in this setting. We predicted gene products with a putative nicotinamide cofactor binding capability through a Rossmann fold domain and monitored the larval development on low and high sugar diets following a ubiquitous knockdown of these candidates (Figure 1A). Knockdown of the candidate genes yielded hits displaying a genotype effect (impaired development irrespective of diet) as well as those displaying a strong enhancement of the phenotype on a high sugar diet (sugar intolerance) (Figure 1B and Supplementary Table 1). Collectively, this implies that nicotinamide cofactor binding proteins have a key role in maintaining homeostasis upon high sugar feeding. One of the hits displaying a strongly sugar-dependent developmental impairment was transcriptional cofactor C-terminal Binding Protein (CtBP) (Figure 1C and Supplementary Figure 1A). Further experiments on larvae carrying a hypomorphic combination of *ctbp* mutant alleles revealed impaired survival on sugar-only diet (Figure 1D and Supplementary Figure 1B-C), showing comparable sensitivity to sucrose, glucose and fructose (Supplementary Figure 1D). The poor survival of CtBP deficient animals on sugar-containing diet was not due to food avoidance behavior, as they displayed increased frequency of mouth hook movements on sugar-only diet (Figure 1E). In contrast to the phenotypes on sugar-containing diet, CtBP deficient larvae survived better on full starvation when compared to control animals (Figure 1F and Supplementary Figure 1E), further emphasizing the sugar-dependency of the CtBP loss-of-function phenotypes.

**Figure 1.**
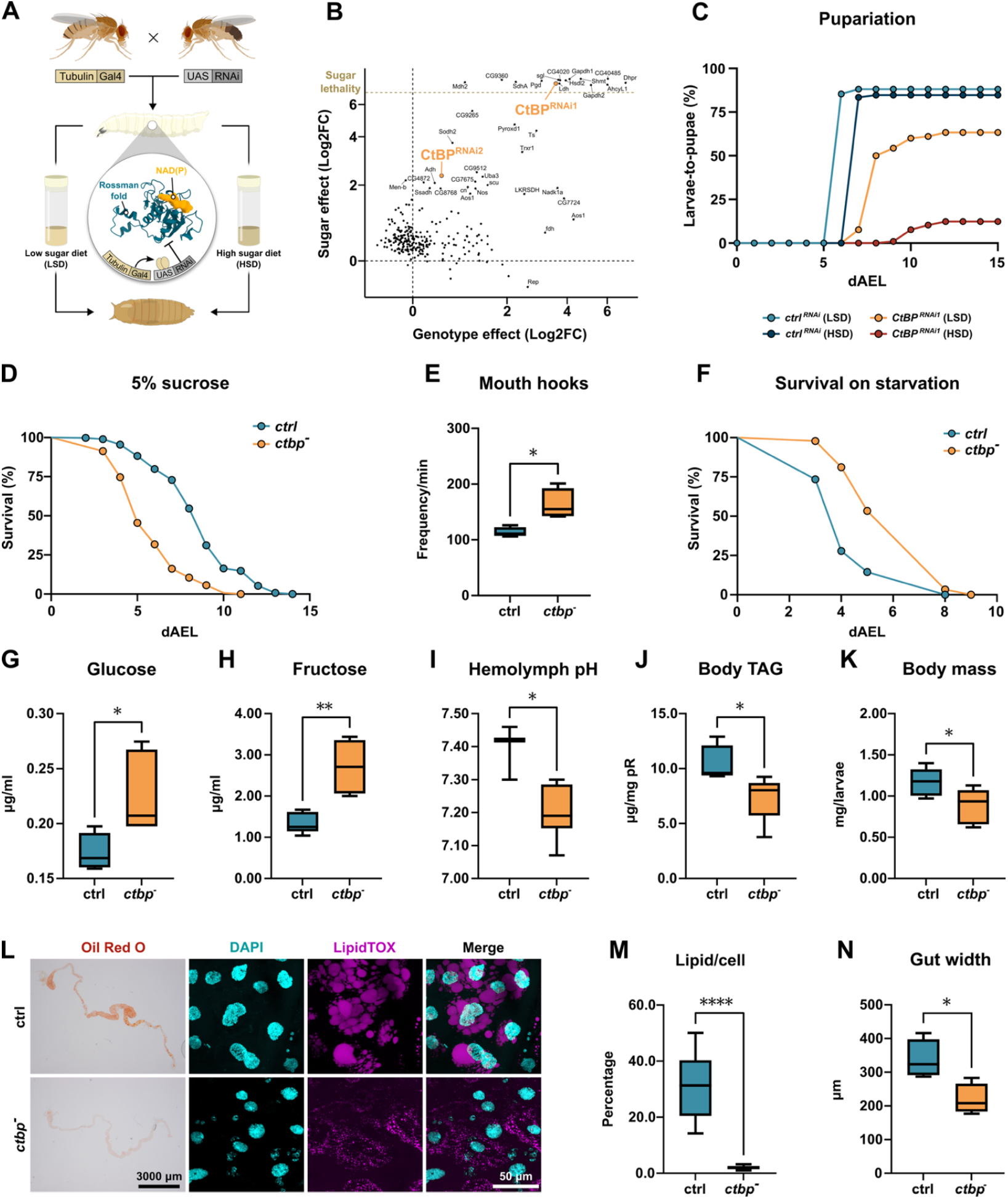
CtBP regulates larval energy metabolism and is necessary for sugar tolerance. **(A)** Schematic overview of the whole-body RNAi screen for sugar intolerance, focusing on proteins with Rossmann fold with predicted nicotinamide cofactor binding activity. **(B)** Genetic screen identifies several genes necessary for sugar tolerance, including *C-terminal binding protein* (*CtBP*). Pupariation index (PI), rewarding for rapid pupariation and high survival rate, was calculated for each condition. X-axis displays the genotype effect, i.e. log_2_ fold change (Log2FC) of PI between control and experimental genotype on low sugar diet (LSD). Y-axis displays the sugar intolerance, i.e. Log2FC of PI of each genotype on LSD vs. high sugar diet (HSD). **(C)** Whole-body knockdown of *CtBP* results in sugar intolerance, manifested as increased lethality and delayed larval development on HSD (20% sucrose). Pupariation rate data for control and *CtBP* RNAi larvae on HSD were compared using the log-rank test (χ^2^ = 109.3, *P* < 0.0001). **(D)** Whole-body *ctbp* loss-of-function (*CtBP*^*03463*^*/CtBP*^*87De-10*^) decreases sugar tolerance compared to heterozygous control (+/*CtBP*^*0346*^). Survival data were compared using the log-rank test (χ^2^ = 351.0, *P* < 0.0001). **(E)** *ctbp*-deficient (*CtBP*^*03463*^*/CtBP*^*87De-10*^) larvae exhibit a hunger-like behavior on a sugar-only diet (5% sucrose), quantified by the number of mouth hook movements per min. Two-tailed t-test, P = 0.0135. **(F)** *ctbp* mutant (*CtBP*^*03463*^*/CtBP*^*87De-10*^) larvae have higher starvation resistance compared to control (+/*CtBP*^*0346*^) (log-rank test, χ^2^ = 50.48, *P* < 0.0001). *ctbp*-deficient (*CtBP*^*03463*^*/CtBP*^*87De-10*^) larvae have elevated levels of glucose **(G)** and fructose **(H)** in their hemolymph on sugar-only diet compared to control (+/*CtBP*^*0346*^) animals (two-tailed t-test, P = 0.0297 and P = 0.0012, respectively). **(I)** Loss of *ctbp* function (*CtBP*^*03463*^*/CtBP*^*87De-10*^) leads to hemolymph acidification (two-tailed t-test, P = 0.0135). This is accompanied by reduced accumulation of triacylglycerols **(J)** and decreased body mass **(K)** (two-tailed t-test, *P* = 0.0177 and *P* = 0.0403, respectively). **(L)** *ctbp*-deficient (*CtBP*^*03463*^*/CtBP*^*87De-10*^) larvae have reduced intestinal lipid content quantified in **(M)** as a lipid volume (mm^3^) per cell (two-tailed t-test, *P* = 0.0403). **(N)** The intestines of *ctbp*-deficient (*CtBP*^*03463*^*/CtBP*^*87De-10*^) larvae are significantly thinner than those of control larvae (+/*CtBP*^*0346*^) (two-tailed t-test, *P* = 0.0175).

To further characterize the physiological role of CtBP, we analyzed the energy metabolism of CtBP-deficient larvae. Consistent with impaired sugar catabolism, the levels of circulating glucose (Figure 1G) and fructose (Figure 1H), were elevated in *ctbp* mutant larvae. Moreover, *ctbp* mutants also displayed hemolymph acidification (Figure 1I), a phenotype earlier associated with defective glucose metabolism^19,21^. Moreover, the relative levels of triacylglycerols were lower in *ctbp* mutant larvae than in controls (Figure 1J), which corresponded with their lower total body mass (Figure 1K). As intestine is a key organ of *Drosophila* lipid biosynthesis^22^, we analyzed the lipid levels in the larval midgut. Staining of neutral lipids with Oil Red O and LipidTOX uncovered a prominent reduction of midgut lipid droplets (Figure 1L-M). Moreover, the midguts of *ctbp* mutants were significantly thinner than those of control larvae (Figure 1N). In sum, CtBP has a major role in the regulation of energy metabolism and growth of *Drosophila* larvae.

### CtBP acts in enteroendocrine cells to control organismal metabolism

How does CtBP control organismal metabolism? To identify possible causes of impaired metabolism in *ctbp* mutants, we performed an RNA sequencing (RNA-seq) profiling of whole larvae deficient for CtBP (Supplementary Figure 2A-D). Notably, enrichment analysis of the RNA-seq data identified deregulation of lipid metabolism, as well as decreased expression of peptide hormones, many of which are produced in the intestine. This, together with the intestinal phenotypes (Figure 1L-N), led us to analyze the midgut-specific role of CtBP more closely. Interestingly, we observed that in animals fed on high sugar diet (HSD) the number of Prospero-positive (Pros^+^) enteroendocrine (EE) was significantly reduced, when compared to control animals (Figure 2A-B). The reduction of EE cell numbers in *ctbp* mutants was not specific to a particular type of EE cells, since numbers of several subpopulations of EE cells, including those expressing diuretic hormone 31 (Dh31), tachykinin (Tk), and allatostatin A (AstA), were reduced in *ctbp* mutant larvae (Figure 2C and D). Consistently, the mRNA levels of the enteroendocrine hormones, produced by these EE cells, were lower in *ctbp* mutants in comparison to control (Supplementary Figure E).

**Figure 2.**
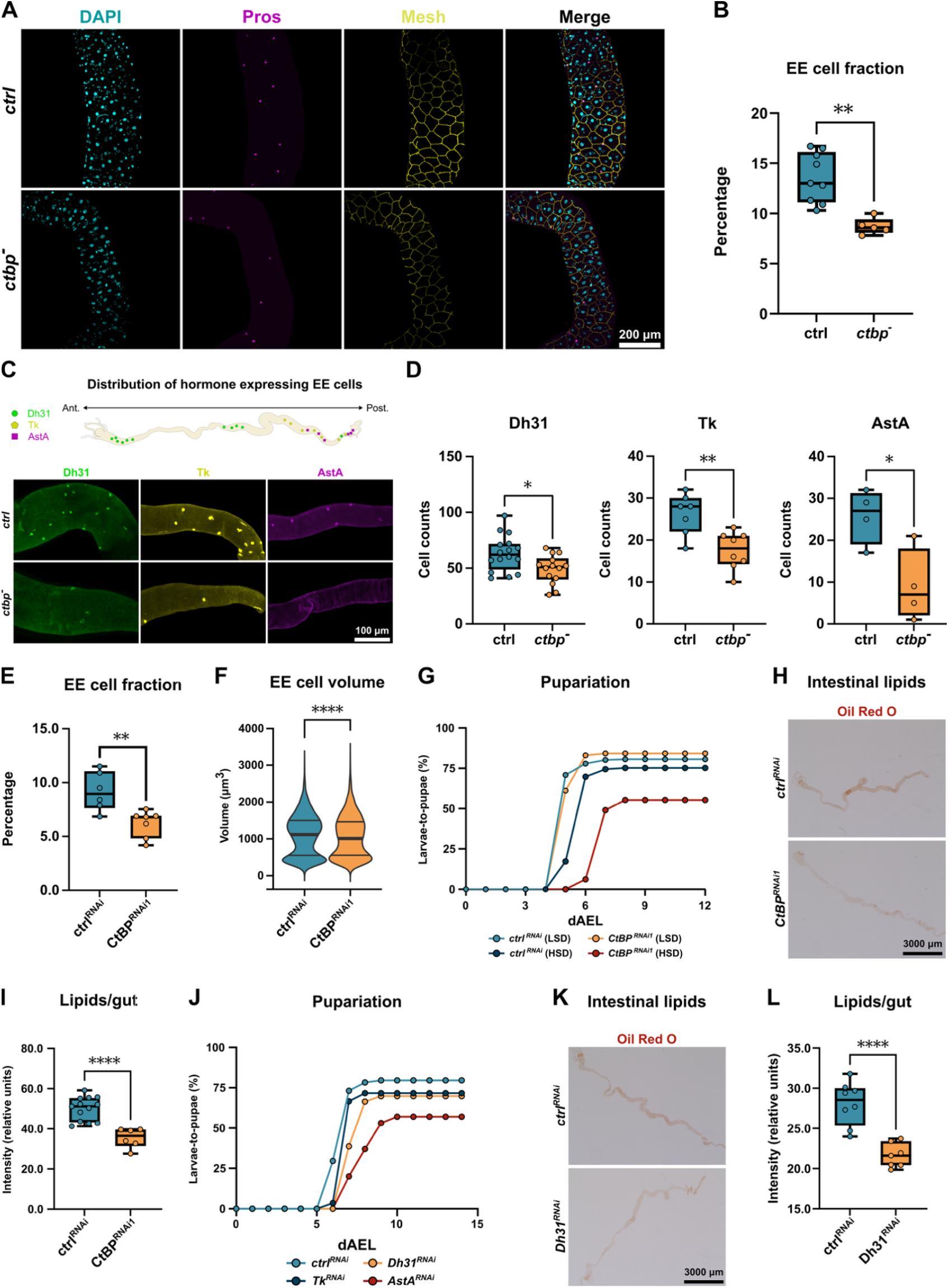
CtBP acts in enteroendocrine cells to control organismal metabolism. **(A)** *ctbp* deficiency (*CtBP*^*03463*^*/CtBP*^*87De-10*^) in *Drosophila* larvae results in a reduction on relative number of EE cells on HSD (30% sucrose), quantified in **(B)**. The data were compared with unpaired t-test, P = 0.0012. **(C)** Upper panel: The distribution of Dh31-, Tk-, and AstA-expressing EE cells in *Drosophila* larvae: a schematic overview. Lower panel: Whole-body *ctbp* loss-of-function (*CtBP*^*03463*^*/CtBP*^*87De-10*^) results in a loss of Dh31-, Tk-, and AstA-expressing EE cells on HSD. **(D)** Quantification of C. Two-tailed t-tests, *P < 0.01; **P = 0.0032. **(E)** Targeted loss of *CtBP* in EE cells (Pros-GAL4>CtBP RNAi) leads to reduced relative number of EE cells in larvae on HSD. Two-tailed t-test, P = 0.0046. **(F)** EE cell specific loss of *CtBP* function (Pros-GAL4 > CtBP^RNAi^) on HSD reduces EE cell size (Mann-Whitney U test, P = 0.00313). **(G)** Loss of *CtBP* function in EE cells (Pros-GAL4>CtBP RNAi) leads to sugar intolerance of *Drosophila* larvae. The pupariation rate of larvae fed on HSD (20% sucrose) were compared with the log-rank test (χ^2^ = 88.6, *P* < 0.0001). **(H)** Intestinal lipid levels are reduced in larvae with EE-specific CtBP deficiency (Pros-GAL4>CtBP RNAi), quantified as relative Oil Red O intensity in **(I)**, unpaired t-test, **\*\*\*\***P < 0.0001. **(J)** Loss of Dh31, Tk and AstA function in EE cells (Pros-GAL4> RNAi) leads to sugar intolerance of *Drosophila* larvae. The pupariation rate of larvae fed on HSD (20% sucrose) were compared using a log-rank test with the adjustment for multiple comparisons (χ^2^ = 59.4, *P* < 0.0001). (**K**) Intestinal lipid levels are reduced in larvae with EE-specific Dh31-deficiency (Pros-GAL4>Dh31 RNAi), quantified as relative Oil Red O intensity in (**L)**, unpaired t-test, **\*\*\*\***P < 0.0001.

Next, we wanted to explore the possibility that CtBP controls EE cells cell-autonomously. In line with this, knockdown of CtBP in EE cells caused phenotypes resembling those observed in the whole-body mutants. The proportion of Pros^+^ EE cells in the larval intestine of animals with EE-cell specific knockdown of CtBP was lower than in controls (Figure 2E). In addition to the reduced cell numbers, also the mean size of Pros^+^ cells was moderately reduced by EE-cell specific knockdown of CtBP on high sugar diet (Figure 2F). The role of CtBP in EE cells was reflected to organismal physiology, as evidenced by impaired sugar tolerance (Figure 2G) and reduced midgut lipid accumulation (Figure 2H, I) upon knockdown in Pros^+^ cells. To test the contribution of individual EE hormones to the observed phenotypes, we analyzed the roles Dh31, AstA, and Tk. Sugar tolerance was strongly impaired by knockdown of AstA and to a lesser extent by that of Dh31 and Tk (Figure 2J), while intestinal lipid storage was significantly reduced by knockdown of Dh31 (Figure 2K, L). Collectively, these data support the role of EE cell specific CtBP function in regulating organismal energy metabolism.

### Nicotinamide redox cofactors control EE cell function and sugar tolerance

Considering the observed sugar regulation of EE cells and the role of CtBP as an NADH binding protein^15^, we hypothesized that nicotinamide cofactors and their redox balance are involved in the regulation of EE cells. To examine whether redox state of EE cells is responsive to dietary sugar intake, we generated transgenic flies expressing genetically encoded NAD^+^/NADH (SoNar) and NADPH/NADP^+^ (iNap) sensors^23,24^ under the control of GAL4/UAS (Figure 3A). To analyze the role of sugar catabolism in EE cell redox balance, we compared the redox balance of larvae fed a high sugar diet vs larvae fed on diet containing 2-deoxy-glucose (2-DG), an inhibitor of glucose catabolism. Diet-derived 2-DG significantly increased NAD^+^/NADH (Figure 3A) and decreased NADPH/NADP^+^ (Figure 3A) ratio, demonstrating that diet-derived sugars regulate EE cell intrinsic redox balance of nicotinamide cofactors.

**Figure 3.**
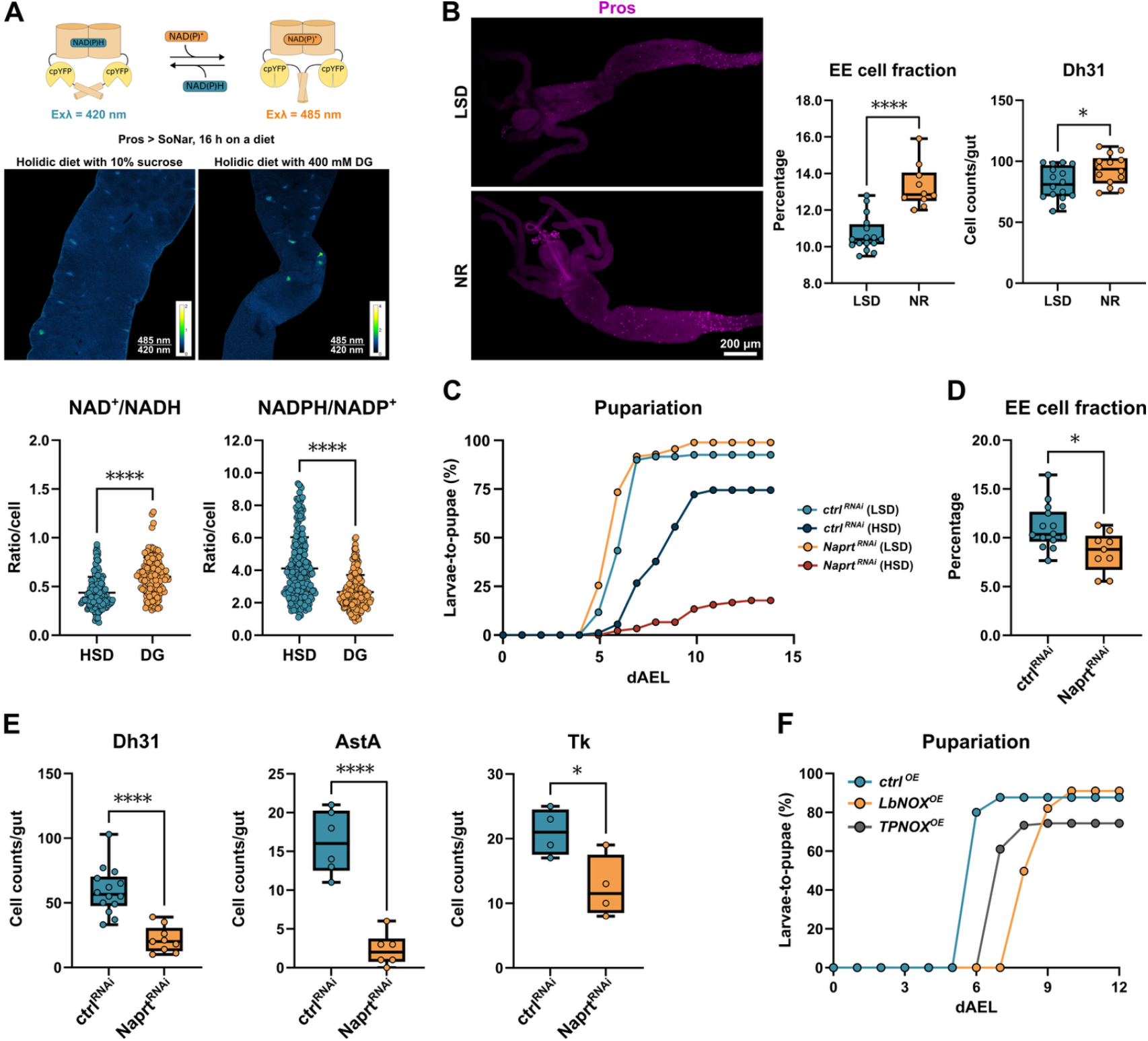
Nicotinamide redox cofactors control EE cell function. **(A)** The ratiometric SoNar and iNap sensors were expressed by Pros-GAL4^ts^ driver to monitor the NAD^+^/NADH and NADPH/NADP^+^ ratios in EE cells. GAL4 expression was activated 16 h prior EE cell redox state measurements. EE cells of HSD fed larvae showed elevated reduction of NAD^+^ and NADP^+^ in EE cells when compared to those of 2-deoxyglucose (2-DG)-fed animals. The data were analyzed using the Mann-Whitney test, with ****P < 0.0001. **(B)** Feeding larvae with nicotinamide riboside (NR), a precursor of nicotinamide adenine dinucleotide, increases the proportion of EE cells (Pros^+^) in the *Drosophila* intestine. Data were analyzed using an unpaired t-test (****P < 0.0001). NR feeding also increases the number of Dh31-positive cells, with statistical significance determined by unpaired t-tests (*P = 0.0149; ***P = 0.0008). **(C)** EE cell specific knockdown (Pros-GAL4) of *Naprt*, a critical enzyme in NAD biosynthesis, leads to sugar intolerance. Pupariation rates of control and experimental genotype on HSD were compared using the log-rank test (χ^2^ = 64.2, *P* < 0.0001). **(D)** EE cell specific knockdown (Pros-GAL4) of *Naprt* reduces relative EE cell numbers in the *Drosophila* intestine as well as **(E)** number of Dh31, AstA and Tk positive cells. The data were analyzed using unpaired t-test, *P < 0.03; ****P < 0.0001. **(F)** Transgenes expressing the bacterial enzymes LbNOX and TPNOX, which oxidize NADH and NADPH in EE cells, respectively, impair larval development on HSD. Pupariation rate is analyzed using the log-rank test, revealing significant effects for both LbNOX (χ^2^ = 43.8, P < 0.0001) and TPNOX (χ^2^ = 54.1, P < 0.0001).

We next asked whether changes in the total pool of nicotinamide redox cofactors have functional relevance for the EE cells. Feeding larvae with nicotinamide riboside (NR), a precursor of NAD^+^ and NADP^+25^, increased the proportion of EE cells in the midgut (Figure 3B) as well as the number of Dh31-positive cells (Figure 3B), implying that NAD(P)^+^ availability increases EE cell abundance in the midgut. Consistent with this, cell-autonomous inhibition of NAD^+^ biosynthesis through knockdown of nicotinate phosphoribosyltransferase (*Naprt*) impaired larval development on a high sugar diet (Figure 3C), reducing the total number as well as several subpopulations of EE cells (Figure 3D-E). To further understand whether changes in the redox balance of nicotinamide cofactors influence the function of EE cells, we generated transgenic flies, which express bacterial enzymes modulating NAD^+^/NADH and NADPH/NADP^+^ ratios, specifically *Lactobacillus brevis* water-forming NADH oxidase (LbNOX)^26^, and TPNOX^27^, a genetically engineered water-forming NADPH oxidase derived from LbNOX. The overexpression of LbNOX and, to a lesser extent, TPNOX in EE cells delayed larval development on HSD consistent with the functional role of NADH and NADPH in EE cells on sugar-fed animals (Figure 3F).

### CtBP senses the metabolic state of EE cells

Given the findings on the functional role of nicotinamide cofactors in EE cells, we wanted to test the possibility whether CtBP responds to changes in EE cell intrinsic NAD(H) and NADP(H) redox balance. As CtBP dimerization is known to be dependent on NADH binding^12,15^, we decided to use bimolecular fluorescence complementation (BiFC) to monitor the status of CtBP in EE cells. In BiFC, two copies of CtBP, fused with fragments of Cerulean and Venus fluorescent proteins, are simultaneously expressed in EE cells. These transgenes produce a functional fluorescent protein when the Cerulean and Venus fragments are brought into close proximity^28^ through CtBP homodimerization (Figure 4A). Consistent with the CtBP dimer formation, we detected a robust fluorescent signal from the Pros^+^ EE cells (Figure 4B). To test the possible roles of NADH and NADPH in CtBP dimerization, we co-expressed the LbNOX and TPNOX transgenes, which oxidize NADH and NADPH, respectively^26,27^ with the CtBP BiFC system. Strikingly, co-expression of both enzymes significantly reduced the relative number of BiFC^+^ EE cells (Figure 4C), providing evidence that oxidization of both NADH and NADPH can inhibit CtBP homodimerization.

**Figure 4.**
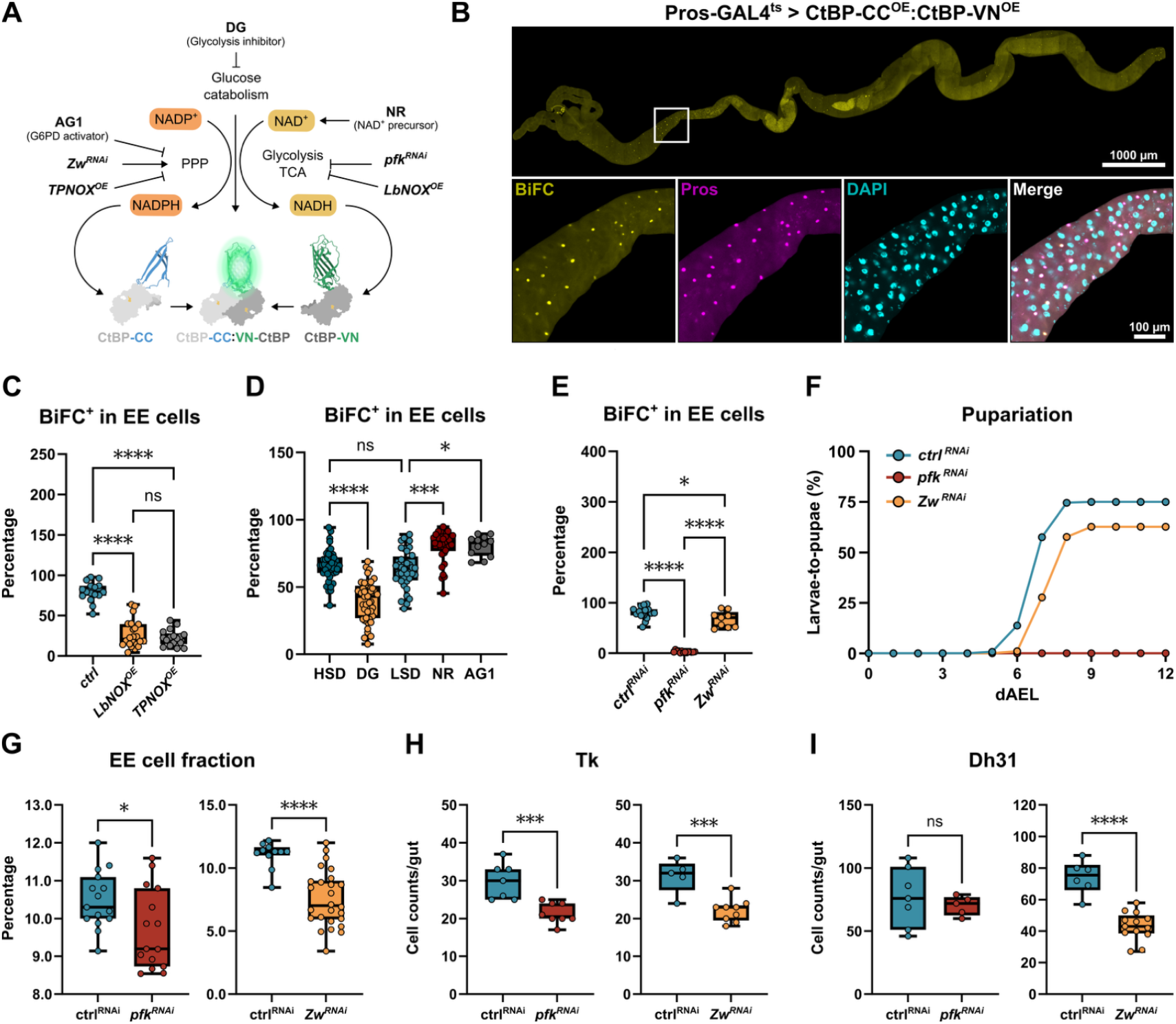
CtBP dimerization is regulated by metabolic pathways affecting NAD(P)H redox balance. **(A)** Schematic presentation of the BiFC system to analyze CtBP homodimerization by the activity of metabolic pathways or enzymes influencing the redox balance of nicotinamide cofactors. GAL4 expression (Pros-GAL4^ts^) was activated 16 h prior measurement for all BiFC experiments. **(B)** CtBP dimerization as measured by BiFC reporters expressed in EE cells (Pros-GAL4^ts^>CtBP-CC^OE^:CtBP-VN^OE^), EE cells marked by anti-Pros antibodies. **(C)** CtBP dimerization (number of BiFC positive cells within EE cell population) is inhibited by LbNOX and TpNOX overexpression, which oxidize NADH and NADPH, respectively. The data are analyzed using ordinary one-way ANOVA, ****P < 0.0001. ^ns^indicates no significant difference in the studied parameter. **(D)** CtBP homodimerization in EE cells (BiFC) is regulated by sugar catabolism and nicotinamide metabolism. A diet containing 2-deoxyglucose (DG) was used to inhibit glucose catabolism. Diets with NR and AG1, a small molecule activator of the enzyme glucose-6-phosphate dehydrogenase (G6PD), were used to modulate the metabolic state. The data are analyzed using one-way ANOVA with Kruskal-Wallis correction for not normal distribution, with *P=0.0143, ***P = 0.0002, ****P < 0.0001. ^ns^indicates no significant difference in the studied parameter. **(E)** Glucose catabolism through glycolysis and the pentose phosphate pathway activates CtBP homodimerization in the EE cells. The enzymes of glycolysis (phosphofructokinase, *pfk*) and the pentose phosphate pathway (G6PD, *Zw*) were depleted in EE cells. The data are analyzed using ordinary one-way ANOVA, with *P=0.0264, ****P < 0.0001. **(F)** Depletion of *pfk* and *Zw* in larval EE cells leads to sugar intolerance. Larval development on HSD (30% sucrose) was fully impaired with *pfk* knockdown (Pros-GAL4>) (χ^2^ = 128.0, P < 0.0001) and modestly delayed by *Zw* knockdown (χ^2^ = 12.1, P = 0.0004), as determined by the log-rank test. **(G)** Knockdown of *pfk* and *Zw* (Pros-GAL4>) in larvae fed on moderate (MSD, 5% sucrose) or high sugar diet (HSD, 30% sucrose) reduces the proportion of EE cells in the larval intestine. These genetic manipulations also decrease populations of Dh31-positive **(H)** and Tk-positive **(I)** cells (unpaired t-test, with **P < 0.01, ***P < 0.001).

This finding prompted us to test the possible role of metabolic pathways influencing the levels and redox balance of nicotinamide cofactors. Dietary supplementation with nicotinamide riboside (NR) significantly increased the relative number of BiFC positive cells within the total EE cell population (Figure 4D), in line with the known role of NADH in CtBP homodimerization^15^. CtBP dimerization was strongly inhibited upon 2-DG treatment (Figure 4D), indicating that CtBP activity in EE cells depends on glucose catabolism. Conversely, supplementation of larval food with AG1, an activator of glucose-6-phosphate dehydrogenase (G6PD)^29^, was sufficient to stimulate CtBP dimerization (Figure 4D), suggesting that activation of the pentose phosphate pathway (PPP), and consequent reduction of NADP^+^ can promote CtBP dimerization. To further test the cell-autonomous roles of glucose catabolic pathways in controlling CtBP dimerization, we inhibited glycolysis and PPP in EE cells, by knocking down phosphofructokinase (*pfk*) and G6PD (*Zw*), respectively. While knockdown of *Zw* moderately reduced CtBP homodimerization in EE cells, the inhibition of glycolysis by depletion of *pfk* displayed near-complete inhibition (Figure 4E).

Finally, we wanted to analyze the role of glucose catabolism in EE cell function. PPP inhibition in EE cells modestly delayed larval development on a high sugar diet (Figure 4F) and reduced the number of EE cells as well as the specific subtypes (Figure 4G-I). Knockdown of *pfk* led to drastic sugar intolerance and full larval lethality on HSD (Figure 4F), implying a major regulatory role for glycolysis in controlling EE cell function. Due to the high lethality, we were not able to analyze the intestines of larvae with EE cell-specific knockdown on a high sugar diet. Depletion of *pfk* under moderate dietary sugar (5% sucrose) reduced the proportion of Pros^+^ EE cells as well as the number of Tk^+^ cells (Figure 4G-I). Considering the strength of the *pfk* loss-of-function phenotype in EE cells, it is conceivable that glycolysis serves other regulatory roles in EE cells, parallel to the regulation of CtBP. Collectively, our data shows that EE cell intrinsic metabolism has a major regulatory role on CtBP dimerization and EE cell function in maintaining organismal sugar tolerance. Our findings support the idea that metabolic changes reflected to the intracellular redox balance of nicotinamide cofactors can be sensed through CtBP homodimerization.

### CtBP target genes regulate EE cell identity and size

How does CtBP exert its control on EE cell function? As CtBP is a transcriptional cofactor, we wanted to identify its genome-wide binding sites in the EE cells, by using targeted DamID, TaDa^30^ (Figure 5A). The TaDa analysis revealed that CtBP associates with >500 target genes in EE cells. Interestingly, the significantly enriched gene groups included pathways related to signaling pathways involved in the control of differentiation and fate (e.g. Notch, Wingless and MAPK) as well as cell size (e.g. mTOR and Hippo) (Figure 5B). Moreover, gene groups related to lipid metabolism were also found enriched as CtBP targets (Supplementary Figure 3A). To better understand the functional roles of CtBP-associated genes, we systematically screened for a selected set of candidate targets, for roles in regulating sugar tolerance and EE cell size. A large fraction of the screened genes, including components of the insulin/mTOR pathway, influenced the size of EE cells (Figure 5C), while having a modest or no effect on sugar intolerance. A smaller subset of genes, with moderate effect on EE cell size, strongly impaired larval sugar tolerance (Figure 5C).

**Figure 5.**
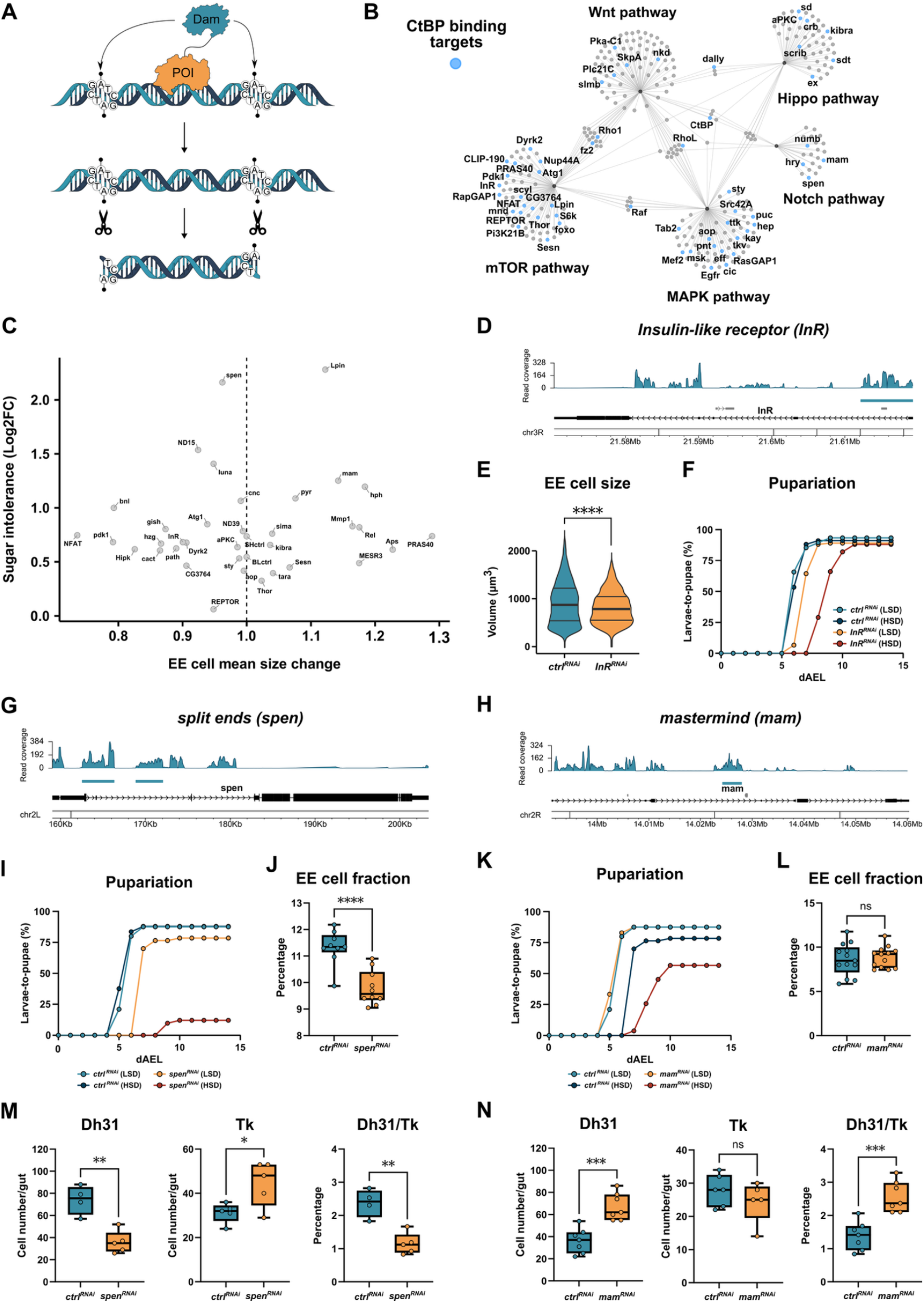
CtBP genomic targets regulate EE cell size and fate. **(A)** Schematic overview presenting the principle of targeted DamID used to identify the genome-wide target loci of chromatin-binding proteins. For determining its genomic binding, CtBP was fused with DNA adenine methyltransferase (Dam) and expressed in EE cells. **(B)** CtBP genomic targets include regulators of fate and growth regulating signaling pathways. CNET plot showing gene overlaps among selected pathways of CtBP targets identified by TaDa. Blue-colored nodes represent CtBP target genes within the pathway network. **(C)** Genetic screen of selected CtBP targets depleted in EE cells (Pros-GAL4>RNAi) reveals regulators of EE cell size and sugar tolerance. The plot shows the effect of the knockdown on EE cell size (X-axis) and sugar intolerance (Y-axis), calculated as the Log2FC of pupariation index of each genotype on LSD vs. HSD (30% sucrose). **(D)** CtBP binding to the genomic locus encoding Insulin-like receptor (*InR*), identified by TaDa. **(E)** Knockdown of InR (Pros-GAL4>RNAi) reduces EE cell size (unpaired t-test, ****P < 0.0001) and **(F)** delays larval development (log-rank test, *χ*^*2*^ = 39.3, *P* < 0.0001) on HSD (30% sucrose). **(G)** Transcriptional regulators Split ends (*spen*) and **(H)** Mastermind *(mam)* are direct targets of CtBP. **(I)** Depletion of *spen* in EE cells (Pros-Gal4>RNAi) results in sugar intolerance. Pupariation rate of larvae with EE-specific *spen* knockdown and its control was analyzed using the log-rank test (χ^2^ = 80.6, *P* < 0.0001). **(J)** Depletion of *spen* in EE cells (Pros-Gal4>RNAi) decreases the proportion of EE cells in *Drosophila* intestine, unpaired *t*-test (****P < 0.0001). **(K)** Knockdown of *mam* in EE cells (Pros-Gal4>RNAi) causes sugar intolerance (log-rank test, *χ*^*2*^ = 64.0, *P* < 0.0001), but **(L)** does not affect the proportion of EE cells in larval intestine, unpaired *t*-test, ^ns^ indicates P > 0.05. **(M)** Knockdown of *spen* in EE cells (Pros-Gal4>RNAi) decreases Dh31 cell numbers and increases Tk cell numbers, lowering the ratio of Dh31/Tk-positive cells (*P = 0.03, **P = 0.0016). **(N)** Knockdown of *mam* in EE cells (Pros-Gal4>RNAi) increases Dh31 cell numbers (***P = 0.001) with no significant effect (^ns^) on Tk-positive EE populations, increasing the ratio of Dh31/Tk-positive cells (***P = 0.001). The data are analyzed with unpaired t-test.

The gene encoding Insulin-like receptor (*InR*) was identified as a CtBP target locus (Figure 5D). The knockdown of *InR* significantly reduced the size of EE cells (Figure 5E) and caused a moderate sugar intolerance (Figure 5F), but did not influence the number of EE cells (Supplementary Figure 3B). Two transcriptional regulators involved in the Notch signaling, namely Split ends (*spen*) and Mastermind (*mam*), were also identified as direct CtBP targets (Figure 5G, H). Knockdown of *spen* caused prominent sugar intolerance, while depletion of *mam* had a more modest effect (Figure 5I, 5K). The knockdown of *spen* significantly reduced the relative number of Pros^+^ EE cells (Figure 5J), while *mam* had no significant influence on Pros^+^ cell numbers (Figure 5L). Surprisingly, both Spen and Mam were found to control the ratio of EE cell subtypes, but in an opposite manner. Loss of *spen* decreased the number of Dh31^+^ positive cells, while increasing the number of Tk^+^ EE cells (Figure 5M). On the other hand, knockdown of the product of *mam* prominently increased the ratio between Dh31^+^ and Tk^+^ EE cells (Figure 5N). In conclusion, our findings show that CtBP is directly associated with genomic loci, which control both EE cell size as well as EE cell identity.

### CtBP associates with Prospero in EE cells

The EE fate-related phenotypes of CtBP, led us to hypothesize possible functional interplay between CtBP and the EE cell fate determinant Pros. To test this hypothesis, we performed targeted DamID analysis on genome-wide Pros targets, which revealed >700 direct target genes (Figure 6A). Comparison between the genome-wide binding profiles of CtBP and Pros revealed that >30% of CtBP targets were common with those of Pros (Figure 6A), implying a significant, but partial, functional overlap. For example, many CtBP target genes in the mTOR and Notch signaling pathways, including the cell fate regulators *spen* and *mam*, were identified as common targets (Figure 6B).

**Figure 6.**
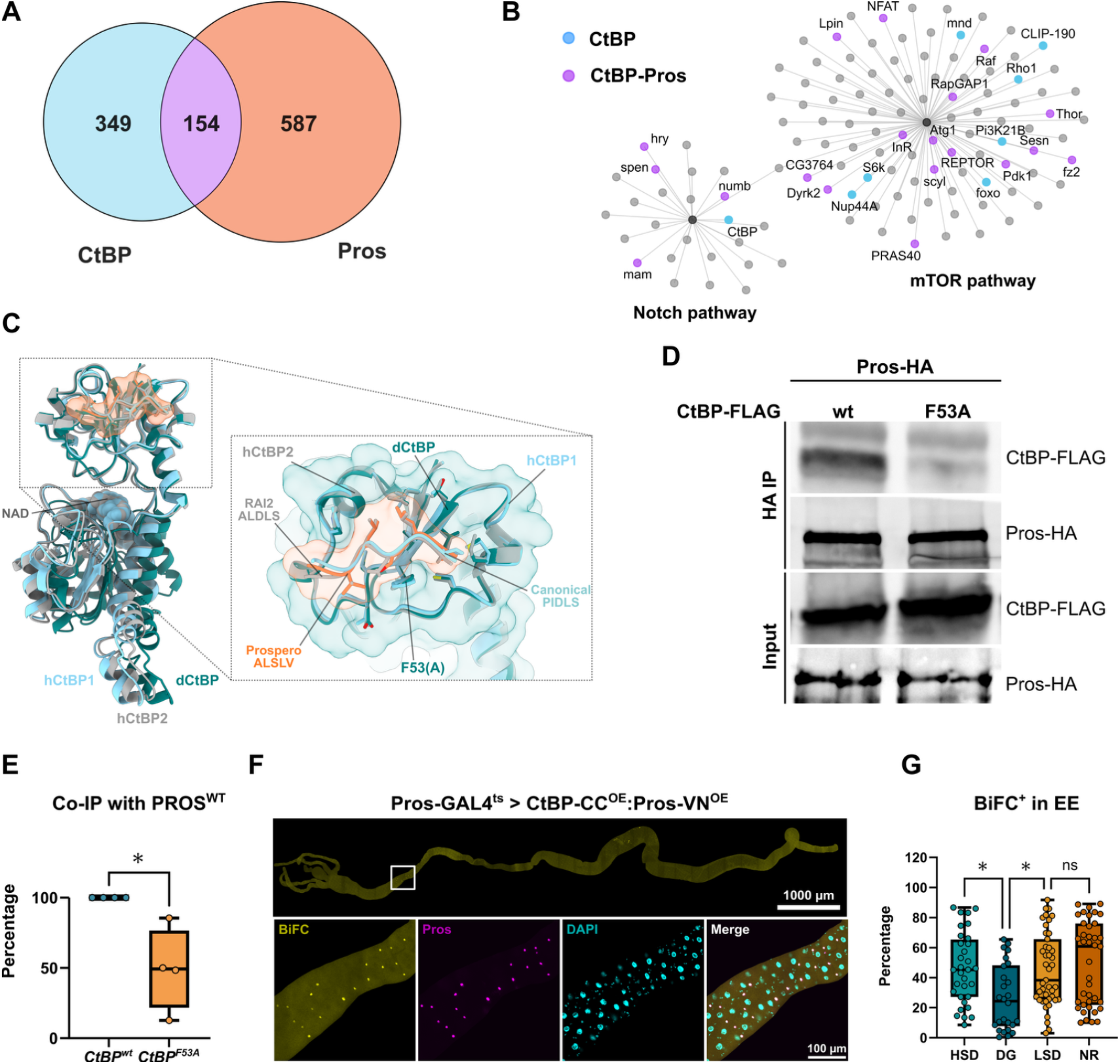
CtBP interacts with Prospero in EE cells. **(A)** CtBP shares >30% of its genomic targets with Pros in EE cells. Comparison of genomic target loci of CtBP and Pros, identified by EE cell-specific TaDa. **(B)** Common targets of CtBP and Prospero include many regulators of the mTOR and Notch pathways. CNET plot showing gene overlaps in the mTOR and Notch pathways among CtBP and Pros targets identified by TaDa. Blue-colored nodes represent CtBP-specific targets, while purple-colored nodes indicate genes co-regulated by CtBP and Pros. **(C)** Conserved substrate-binding cleft mediates interaction between *Drosophila* CtBP (dCtBP) and Prospero. AlphaFold2 multimer model of dCtBP (teal) bound to the Prospero ALSLV motif (residues 1284–1288, orange) is shown superimposed with crystal structures of human CtBP1 (hCtBP1, grey) bound to a canonical PIDLS motif (beige, PDB ID: 1HL3; Nardini et al., 2003), and human CtBP2 (hCtBP2, cyan) bound to a non-canonical ALDLS motif from RAI2 (PDB ID: 8ATI, cyan, Goradia et al., 2024). The zoomed-out view displays the structural alignment of substrate-binding domains across dCtBP, hCtBP1, and hCtBP2, highlighting motif engagement within the conserved PxDLS-binding cleft. The zoomed-in view emphasizes the structural conservation of the CtBP binding pocket and the similar configurations adopted by ALSLV, PIDLS, and ALDLS motifs, despite sequence differences. A computational estimate of the change in free binding energy (ΔΔG) identifies a phenylalanine-to-alanine mutation at position 53 in dCtBP (CtBPF53A) as a key substitution predicted to significantly reduce binding (see also Supplementary Figure 3C). Proteins are shown in cartoon representation, interacting residues as sticks, and NAD as spheres. **(D)** Co-immunoprecipitation of ectopically expressed FLAG-CtBP (CtBP^WT^) and HA-Pros (Pros^WT^) in *Drosophila* S2 cells confirms the ability of CtBP to interact with Pros. The CtBP-Pros interaction is significantly reduced by the CtBP F53A mutation, consistent with structural predictions. **(E)** Quantification of the data is displayed as a percentage of Co-IP relative to CtBP^WT^ and normalized to the input (unpaired t-test *P < 0.03). **(F)** CtBP interacts with Pros in EE cells as revealed by the Bimolecular Fluorescence Complementation (BiFC) assay. **(G)** CtBP-Pros interaction is significantly inhibited by 2-DG feeding, but in contrast to CtBP-CtBP interaction, it remains unaffected by NR feeding. Data are analyzed using the one-way ANOVA with Kruskal-Wallis correction for not normal distribution, with *P < 0.02. ns indicates no significant difference in the studied parameter.

To investigate the potential interaction between *Drosophila* CtBP and Pros, we utilized the AlphaFold2 multimer model^31,32^ incorporating a fragmentation strategy^33^ and augmented confidence metrics^34^. This approach predicted with high confidence (average pLDDT of ∼91, ipTM ∼0.92, and actifpTM ∼0.94), that the Pros motif ALSLV (residues 1284-1288) binds to the cleft at the dCtBP substrate-binding domain (Figure 6C). The binding cleft is conserved across *Drosophila* CtBP and mammalian CtBP1/2 and is a known interaction site for multiple transcription factors and co-regulators via the canonical PxDLS motif^35,36^. Interestingly, recent structural data revealed mammalian CtBP2 bound to RAI2 via a non-canonical ALDLS motif at the same cleft^37^. Superimposing our predicted CtBP-Pros ALSLV-bound conformation with crystal structures of CtBP bound to canonical (PIDLS) and non-canonical (ALDLS) motifs revealed a high degree of structural similarity (Figure 6C). Additionally, AlphaFold2 multimer predicted a similar binding configuration between human CtBP1/2 and the PROX-1 ALPLV motif (residues 438-441), suggesting a potential conservation of the interaction between these proteins in mammals.

To validate the predicted *Drosophila* CtBP-Pros direct binding, we performed co-immunoprecipitation of ectopically expressed FLAG-CtBP and HA-Pros in *Drosophila* S2 cells, combined with *in silico* alanine scanning of the CtBP cleft with the bound Pros motif. Co-immunoprecipitation confirmed CtBP-Pros binding and its disruption by the CtBP F53A mutation (Figure 6D-E), which had been computationally predicted to impair CtBP-Pros binding (Supplementary Figure 3C). To analyze the possible interaction between CtBP and Pros in EE cells, we used the BiFC assay with CtBP fused to a Cerulean fragment and Pros fused to a Venus fragment. The BiFC analysis confirmed the AlphaFold2 prediction and co-IP evidence for CtBP-Pros interaction, as co-expression of CtBP and Pros BiFC fusion proteins in EE cells produced a robust fluorescent signal (Figure 6F), which was not detected in the negative control co-expressing BiFC fusions of Esg and CtBP (Supplementary Figure 3D). We wanted to test the possibility, whether the interaction between CtBP and Pros is nutrient-dependent and observed that feeding of 2-DG, in comparison to a high sugar diet, moderately but significantly, reduced the interaction between CtBP and Pros in EE cells (Figure 6G). However, feeding of NR had no influence on the CtBP-Pros interaction, implying that CtBP can associate with Pros in both monomeric and dimeric forms, similar to what has been observed with other transcription factors^16^. Collectively our data shows that CtBP interacts with EE cell fate determinant Pros to target genomic loci involved in the regulation of EE cell fate.

## DISCUSSION

In this study, we uncover a molecular mechanism, mediated through transcriptional cofactor CtBP, which acts in the interface between nutrient sensing and the control of enteroendocrine (EE) cell identity and function. Through its activity in EE cells, CtBP controls organismal energy metabolism and tolerance to a high sugar diet, underlining the key role for EE cell function in mediating physiological adaptation to dietary sugars. We observed that the EE cell NAD(H) and NADP(H) redox balance is regulated in response to glucose catabolism and that these changes are sensed through the dynamic regulation of CtBP dimerization. Inhibition of glucose catabolism through both glycolysis and the pentose phosphate pathway was sufficient to inhibit CtBP dimerization and similar effects were observed when NADH and NADPH were oxidized through overexpression of bacterially derived *Lb*NOX and TPNOX enzymes. Thus, our findings are consistent with a model, where EE cell-autonomous glucose catabolism, through its influence on redox balance of nicotinamide cofactors, dynamically controls CtBP activity and consequently enteroendocrine function in response to dietary sugar intake. Mechanistically, CtBP interacts with the transcription factor Prospero (Pros), the master regulator of EE cell fate. In genome wide comparison, CtBP binding is observed in a significant subset of Pros targets, including regulators and components of the Notch and insulin/mTOR pathway, contributing to the regulation of EE cell fate and size, respectively. Our genome-wide data on CtBP and Pros chromatin binding is consistent with a model that CtBP is a context-dependent modulator of Pros and that CtBP may also have Pros-independent functions in EE cell regulation.

Our study reveals that enteroendocrine (EE) cell signaling has a major role in regulating organismal adaptation to a high sugar diet, likely through multiple EE hormones. Previous studies in adult *Drosophila* have implicated Neuropeptide F and Bursicon α in sugar-induced metabolic regulation, which control organismal metabolism by modulating the insulin-glucagon/Akh signaling^5,6^. Our study reveals additional EE hormones involved in regulating metabolism in response to dietary sugars. In particular, we observed that Allatostatin A (AstA), earlier identified as negative regulator of lipid storage^38^, displayed a strong contribution to larval sugar tolerance. These findings imply a broader role for EE hormone signaling in sugar metabolism than previously anticipated and call for a systematic comparative analysis to elucidate the functional roles and interactions of all EE hormones in this setting. Previous studies have established the critical role of the sugar-sensing transcription factor complex Mondo-Mlx in maintaining larval viability under high sugar conditions^39^ via transcriptional regulation of metabolic genes in the fat body and intestinal enterocytes^20,40^. The findings uncovering the role of EE cell signaling in sugar tolerance raise the intriguing possibility that the EE cell signaling might display previously unknown functional crosstalk with Mondo-Mlx-mediated intracellular sugar sensing, which should be addressed in future studies.

We also provide evidence for the role of CtBP as a physiologically relevant, redox state-responsive molecular switch. Earlier biochemical studies have shown that CtBP undergoes NADH-mediated dimerization^15^ enabling it to assemble distinct protein complexes involved in gene regulation^15,16,41^. However, whether the NADH-mediated dimerization allows CtBP to sense physiologically relevant changes in cellular metabolism and redox state has remained debatable. Our observations on glycolysis-driven regulation of CtBP dimerization in EE cells align with the existing biochemical reports showing high-affinity binding of NADH by CtBP^12,15^. However, the observed regulation of CtBP dimerization by the pentose phosphate pathway (PPP) activity and NADP(H) redox status is somewhat surprising, given that NADH-binding domain of *Drosophila* CtBP contains a conserved GXGXXG motif, characteristic for proteins binding NADH as a cofactor^41^. While it remains a possibility that high intracellular NADPH allows direct regulation of CtBP dimerization, it is also possible that the observed *in vivo* regulation of CtBP dimerization is due to secondary effects on EE cell metabolism. In addition to impaired NADP^+^ reduction, inhibition of PPP negatively impacts NAD-biosynthesis by limiting the supply of phosphoribosyl pyrophosphate (PRPP)^42^, a precursor of nicotinamide nucleotides via *de novo* and Preis-Handler pathway. Since many cellular processes consume NAD^+^, this may hinder maintenance of soluble NAD(H) levels. Moreover, NADPH is a primary feedback-inhibitor of NAD-kinase, and its depletion can promote the phosphorylation of NAD^+^ to NADP^+^ eventually depleting the NADH pool^43^. Despite the remaining questions regarding the biochemical details of CtBP regulation in EE cells, our data strongly support the view that CtBP dimerization is highly responsive to metabolic changes observed in physiological setting, allowing it to contribute to hormonal regulation of animal physiology in a nutrient responsive manner.

Similar to our observations on CtBP, the mammalian ortholog of *Drosophila* Mondo, ChREBP, was recently shown to be activated by increased reduction of NAD^+^ in hepatic tissue^44^. Collectively, these findings indicate that the molecular sensors for intracellular sugars use the dynamic changes in nicotinamide cofactor redox states as a proxy for cellular metabolic state, in parallel to direct sensing of sugar derivatives^45^. Our genetic screen on Rossmann fold containing proteins contributing to sugar tolerance serves as a valuable resource for future efforts to understand the global role of nicotinamide cofactor-binding proteins in sugar-responsive metabolic adaptation, including Mondo/ChREBP mediated sugar sensing. Our findings on the EE cell specific metabolic role of CtBP extend earlier *in vivo* evidence for CtBP as a modulator of glucose metabolism in the liver^18^. Thus, CtBP appears to have several distinct tissue-specific roles in metabolic regulation, similar to its multifaceted role as a developmental regulator^46^.

Our study uncovers an interaction between CtBP and the EE cell fate regulator Pros, which is mediated through a non-canonical, but conserved, binding motif. Although this interaction was previously observed in a yeast two-hybrid screen^47^, our data provides a biologically relevant context for CtBP-Pros interaction and reveals insight into its structural basis. While we observed a significant overlap between CtBP and Pros genomic targets, it is important to note that both factors also associate independently with many target loci. Shared targets of CtBP and Pros include regulators of EE cell size, which is consistent with the observation that CtBP-deficient animals have smaller EE cells on a high sugar diet. Since cell size regulation is closely linked to anabolic processes, such as translation and secretion^48^, the dynamic control of EE cell size may be relevant in adjusting the secretory capacity of EE cells. Other common downstream targets include *spen* and *mam*, both linked to Notch signaling^49-51^. Previous evidence from adult *Drosophila* intestinal stem cells shows that inhibition of Spen leads to stem cell accumulation and Delta expression, possibly due to a defect in Delta trafficking^52^. Our findings imply that Spen also has a necessary role in maintaining the EE cell identity. Mastermind, a transcriptional coactivator, is a well-established mediator of Notch-induced gene expression^50,51^. Interestingly, our data show that while both genes contribute to EE cell signaling in sugar-fed animals, they regulate specific EE cell subtypes in an opposite manner. This suggests a nuanced role for CtBP in adaptive nutrient-responsive regulation of EE cells, possibly allowing dynamic adjustment of EE cell subtype composition in response to ectopic cues. Future studies addressing the molecular targets of Spen and Mam are likely to yield new insight into the mechanisms regulating the subtype composition of the intestinal EE cell pool.

In conclusion, the results reported here outline a modality that allows the plasticity of the enteroendocrine system and consequent regulation of organismal metabolism through nutrient responsive control of EE cell fate and size. The key molecular mediator of the underlying mechanism is the pleiotropic transcriptional cofactor, CtBP, which uses glucose catabolism-induced changes in cellular redox balance as a proxy for the availability of dietary sugars. Through its interaction with the EE cell lineage specific transcription factor Prospero, CtBP mediates the nutrient-derived information to the gene regulatory network responsible for the control of EE cell identity and size. Considering the biomedical importance of EE cell signaling^2^, the molecular mechanisms discovered here may open new directions for further research to therapeutically modify the abundance, secretory capacity and subtype composition of the EE cell pool.

## MATERIALS AND METHODS

### *Drosophila* lines and husbandry

*Drosophila* stocks were kept under controlled conditions (25°C with a consistent 12 h light-dark circadian cycle), fed a basic laboratory diet composed of 6.5% malt (w/v), 3.2% semolina (w/v), 1.8% dry baker’s yeast (w/v), 0.6% agar (w/v), 0.7% propionic acid (v/v), and 2.5% nipagine (methyl paraben) (v/v).

The experiments with *ctbp* mutants presented in the main figures were done using allelic combinations of *CtBP*^*03463*^ (BDSC: 11590) and *CtBP*^*87De-10*^ (KDSC: 106438) earlier characterized as a hypomorph and a null mutation, respectively^53^. These alleles were isogenized with *w*^*1118*^ line (BDSC: 6326), which was used as control in combination with *CtBP*^*03463*^. Sugar sensitivity of mutant alleles *CtBP*^*KG07519*^ (KDSC: 111116) and *CtBP*^*GS15006*^ (KDSC: 206039) was also tested. The Gal4 drivers used in this study: *Tub-Gal4* (BDSC: 5138); *Ubi-Gal4* (BDSC: 32551); *Tub-Gal80*^*ts*^, *Pros-Gal4*^54^ referred also as *Pros-Gal4*^*ts*^, *NPF-Gal4* (BDSC: 25681), *Dh31-Gal4* (BDSC: 51988). The transgenes and RNAi lines from publicly available sources used in the study: *UAS-CtBP*^*RNAi1*^ (BDSC: 32889); *UAS-CtBP*^*RNAi2*^ (VDRC: 107313); *UAS-Luc*^*RNAi*^ (BDSC: 31603, used as a control for TRIP lines); *UAS-w*^*1118*^ (VDRC: 60100, used as a control for the VDRC lines); *UAS-spen*^*RNAi*^ (BDSC: 33398); *UAS-mam*^*RNAi*^ (BDSC: 28046); *UAS-InR*^*RNAi*^ (VDRC: 992); *UAS-Zw*^*RNAi*^ (VDRC: 101507); *UAS-pfk*^*RNAi*^ (BDSC: 34336); *UAS-Dh31*^*RNAi*^ (BDSC: 41957); *UAS-Tk*^*RNAi*^ (BDSC: 25800); *UAS-AstA*^*RNAi*^ (BDSC: 25866); *UAS-Naprt*^*RNAi*^ (BDSC: 62264); *UAS-CtBP-ORF-CC* (FlyORF: F004803); *UAS-CtBP-ORF-VN* (FlyORF: F004799); *UAS-Pros-ORF-VN* (FlyORF: F004799); *UAS-Esg-ORF-CC* (FlyORF: F003375). The list of lines used in RNAi screens is presented in Supplementary Tables 1 (whole-body RNAi screen) and 2 (EE specific RNAi screen). Transgenic lines generated in this study: *UAS-Luc, UAS-LbNOX, UAS-TPNOX, UAS-SoNar, UAS-iNap, UAS-SoNar(iNap) control, UAS-DAM-CtBP; UAS-DAM-Pros; UAS-DAM-control*.

### Generation of transgenic lines

The sequences of CtBP (isoform A) and Prospero (isoform H) were generated by gene synthesis (Azenta Life Sciences) and inserted into the pUAST-attB-LT3-NDam vector (a kind gift from A. Brand laboratory) using 5’ XhoI and 3’ XbaI restriction sites. Sequence-verified vector was then introduced into the yw fly strain at the attP2 site using the PhiC31 integrase system. Additionally, a control fly line containing an unconjugated Dam construct was also created. The FLAG-tagged sequences of LbNOX and TPNOX, were codon optimized for expression in the *Drosophila* model using Azenta Life Science services and inserted into pUAST vector using 5’EcoRI and 3’XhoI restriction sites. The sequences for the NAD^+^/NADH redox sensor SoNar, the NADPH/NADP^+^ redox sensor iNap, and their redox-insensitive mutant control were generated by gene synthesis and cloned into the pUAST vector using 5′ EcoRI and 3′ KpnI restriction sites. Sequence-verified vectors of LbNOX and TPNOX were inserted into the w strain at Zh51C site, whereas SoNar and iNap vectors as well as their control vector were inserted into the VK0037 landing site.

### Generation of plasmids

To generate an HA-tagged Prospero construct, the coding sequence of pros-PM was cloned upstream of tandem HA epitopes into the pUAST vector using 5’ Bgl II and 3’ Xho I restriction sites. For the FLAG-tagged CtBP construct, the CtBP-RA open reading frame was fused via a GGS_3_ linker to a C-terminal 3×FLAG tag and inserted into pUAST using the same restriction sites. The CtBP^F53A^ variant was generated by substituting phenylalanine 53 with alanine. All plasmids were generated by gene synthesis using Azenta Life Sciences (US) services.

### Experimental diets

All experimental diets contained 0.75% agar (w/v), 0.7% propionic acid (v/v), and 2.5% nipagine (v/v). Low sugar diets (LSD) included 10% or 15% dry baker’s yeast (Lowan Whole Foods, w/v) as the primary nutrient source. High sugar diets (HSD) in addition to the yeast, had 15, 20 or 30% sucrose (Dansukker, w/v). The concentration of sucrose used in specific experiments is detailed in the corresponding figure legends. For the experiments on intracellular nicotinamide redox ratios, HSD stands for *Drosophila* larvae-optimized chemically defined diet^55^ supplemented with 10% sucrose. Sugar-only diet contained 5% sucrose or 15% sucrose (Sigma-Aldrich, S9378), fructose (Sigma-Aldrich, F0127) or glucose (Sigma-Aldrich, G7021). The starvation diet consisted of agarized water without any nutrients. NR and AG1 diet contained yeast in concentration of 15% (w/v) and 500 µM nicotinamide riboside chloride (Sigma-Aldrich, SMB00907) or 10 µM glucose-6-phosphate dehydrogenase activator AG1 (MedChemExpress, HY-123962). DG diet included 400 mM 2-deoxy-D-glucose (Acros Organics, AC111980050) in addition to 15% yeast (w/v) or the chemically defined diet. Two experimental feeding strategies were employed in this study: long-term and acute treatments. For long-term dietary treatments, carefully staged first instar larvae, reared on yeast-supplemented apple juice plates (33.3% apple juice, 1.75% agar, 2.5% sugar, 2.0% nipagin) at 25°C, were transferred to vials containing the experimental diets. Activation of the *Pros*^*ts*^ driver was initiated at this point, when applicable. For acute treatments, larvae were initially reared on a low-sugar diet (LSD) or standard diet and then transferred as late second instar larvae to experimental diets for 16 hours with simultaneous activation of Pros^ts^ driver.

### RNAi screens, larval survival and quantification of sugar intolerance

For the whole-body RNAi screen, gene silencing was induced using the ubiquitously expressed *Tubulin-Gal4 (Tub-Gal4)* driver. In cases where larval lethality occurred irrespective of diet, the *Ubiquitin-Gal4 (Ubi-Gal4)* driver was used instead. For EE cell-specific gene silencing, a temperature-sensitive *Prospero-Gal4 (Pros-Gal4*^*ts*^) driver was used. For the RNAi screens, thirty carefully staged first instar larvae were transferred into vials containing either a low (LSD) or high sugar diet (HSD). To assess survival of *ctbp* mutants, larvae were transferred to vials containing either a sugar-only diet (5% or 15% sugar) or subjected to complete starvation. Brilliant Blue FCF dye (Sigma-Aldrich, 861146) was added to the diet to help visualize dead larvae. Pupariation and survival were monitored every 24 hours. Developmental fitness was quantified by assessing the effects of diet and genotype on pupariation. The diet effect was calculated as the log_2_ fold change in pupariation index for larvae of the same genotype on LSD versus HSD. The genotype effect was calculated as the log_2_ fold change in pupariation index between experimental and control genotypes, independent of diet. Pupariation index was calculated as in^56^

### Metabolites and pH measurements

Hemolymph and whole-body glucose levels, as well as triacylglycerol content, were measured in 3^rd^ instar larvae as previously described^19^. Hemolymph fructose was quantified using a modified Seliwanoff’s method^57^, and hemolymph pH was assessed using pyranine dye, following established protocols^19,21^.

### Mouth hook assay

For the mouth hook assay, larvae were reared on standard laboratory food until the early 3^rd^ instar stage. A single larva was then transferred to a plate containing food supplemented with 5% sucrose and food dye Brilliant Blue FCF (FD&C Blue No. 1). After a 3-hour preincubation period, the larva was recorded for 1 minute, and mouth hook movements were quantified.

### Immunohistochemistry

For immunofluorescence staining, intestines were dissected in phosphate-buffered saline (PBS), and then fixed in 4% paraformaldehyde overnight at 4 °C. Fixed guts were washed with PBS containing 0.1% Triton X-100 and blocked using 1% bovine serum albumin (BSA) for one hour. Following blocking, the tissues were incubated with antibodies against Prospero (1:500, Developmental Studies Hybridoma Bank, MR1A), Mesh (1:1000, a kind gift from Mikio Furuse), AstA (Developmental Studies Hybridoma Bank, 5F10C), Dh31 or Tk (a kind gift from Jan Veenstra). Fluorescent secondary antibodies were obtained from Invitrogen (Thermo Fisher Scientific). Vectashield mounting medium with DAPI (Vector Laboratories, 1200-10) was used to stain DNA.

### Lipid analysis

For lipid staining, midguts from third instar larvae were dissected in PBS and fixed in 4% paraformaldehyde for 30 min, then washed thrice with PBS. The guts were then stained either with LipidTOX (H34477, Thermo Fisher Scientific), diluted in PBS at a ratio 1:400 for 20 min or with 60% Oil Red O for 10 min. Prior to Oil Red O staining, the guts were preincubated with 60% isopropanol for 5 min. After staining, the midguts were washed again thrice with PBS and mounted using Vectashield Mounting Medium with DAPI (Mediq) or 70% glycerol.

### Microscopy

Fixed and immunostained *Drosophila* larval intestines were mounted between a microscope slide and a coverslip using 60 μm spacers. For visualization of neutral lipids stained with Oil Red O, samples were imaged using a Leica DMS1000 B light microscope. Imaging of immunostained midguts was performed using an Aurox Clarity spinning disk confocal microscope, and image processing was carried out with ImageJ software. High-resolution imaging of LipidTOX-stained samples and live imaging of iNAP and SoNar redox sensors was conducted using a Leica TCS SP8 STED 3X CW 3D microscope. Signal intensities were quantified as mean voxel or pixel values within segmented regions of interest.

### Quantification of enteroendocrine cell size and number

Confocal image segmentation and enteroendocrine (EE) cell quantification were performed semi-automatically using IMARIS software (versions 9.3– 10.2). EE cell size was quantified using a custom-made automated script based on nuclear segmentation from anti-Pros antibody staining.

### Measurement of NAD^+^/NADH and NADPH/NADP^+^ redox ratios

EE specific changes in NAD^+^/NADH and NADPH/NADP^+^ redox ratios were identified using genetically encoded SoNar and iNap sensors under the control of GAL4/UAS. They utilize NAD(P)H binding domain of bacterial T-Rex fused to cpYFP^23,24^ and display differential fluorescence upon binding of reduced (485 nm) vs. oxidized (420 nm) cofactor at a single emission at 520 nm, allowing monitoring of NAD^+^/NADH and NADPH/NADP^+^ ratio *in vivo*. For the measurements, 3^rd^ instar larvae were dissected in *Drosophila* Schneider’s medium after 16 hours of growth on either a chemically defined diet^55^ supplemented with 10% sucrose (HSD) or with 2-deoxy-D-glucose (DG) and used for live imaging. NAD^+^/NADH and NADPH/NADP^+^ ratios were quantified as the ratio of fluorescence intensities for NAD(P)^+^ or NAD(P)H-bound sensors in manually segmented enteroendocrine (EE) cells.

### *Drosophila* cell culture

S2 cells (ATCC, CRL-1963) were grown at 26°C in Scheider’s medium (Thermo Fisher Scientific; 21720024) supplemented with 10% heat-inactivated FBS (Merck, 10270106) and 50 µg/ml streptomycin. S2 cells were transfected by Effectene kit (Qiagen, 301427) with pAct-Gal4 together with selected pUASTattB-based plasmids.

### Co-Immunoprecipitation (Co-IP) assay and Western blotting

For the co-immunoprecipitation, S2 cells were collected and homogenized in ice-cold lysis buffer (20 mM Tris-HCl, pH 7.5, 150 mM NaCl, 1 mM EDTA, 1% Triton, protease inhibitor cocktail tablet and PhosSTOP tablet). The lysed cells were centrifuged (1300 g, 4 min, 4°C) and the protein concentration was determined using a BCA assay. A total of 1.2 mg of lysate was incubated with 10 µl of anti-HA magnetic beads (Thermo Fisher Scientific, 88836) for 2 hours at 4°C under agitation. The beads were collected magnetically and washed three times with lysis buffer. Proteins were eluted from the beads with 15 µl of Laemmli buffer.

For Western blotting, S2 cells were collected and homogenized in the same buffer as described for the co-immunoprecipitation assay. Primary mouse anti-FLAG (1: 2500, Sigma-Aldrich, clone M2) and rabbit anti-HA (1:2000, Abcam, ab9110) were used to bind CtBP and Prospero tagged with FLAG and HA, respectively. The fluorophore-conjugated secondary antibodies StarBright Blue 700 anti-rabbit (1:2500, BioRad) and StarBright Blue 520 anti-mouse (1:2500, BioRad) were used. The ChemiDoc XRS+ system (BioRad) was used to image the bands. To assess co-immunoprecipitation efficiency, the ratio of co-IP to IP was quantified. Co-IP levels were expressed as a percentage relative to CtBP^WT^ and normalized to the input.

### RNA sequencing and tissue specific expression analysis

Carefully staged 1^st^ instar larvae were collected 24 hours after egg laying and kept on plates with a regular lab food at a set density. At 48 hours after egg laying, early 2^nd^ instar larvae were transferred to plates containing sugar-only diet (5%) with the blue food dye. After 8 hours of diet exposure, 30 actively feeding larvae per sample were collected and immediately frozen in liquid nitrogen. The larvae were homogenised and total RNA was extracted using the Nucleospin RNA II kit (Macherey-Nagel) following the manufacturer’s guidelines. RNA libraries were prepared using the TruSeq Stranded mRNA kit (Illumina), and sequenced on the Illumina NextSeq500 system to achieve an average depth of 20 million reads per sample, utilizing the 75bp cycle kit (Illumina).

The quality of the raw sequencing data was evaluated using FastQC^58^ version 0.11.2. Reads were trimmed using Trimmomatic version 0.33^59^, ensuring each was at least 36 bases long and met quality scores of 15 per base and 20 across both strands with a 4-base sliding window. Reads were aligned to the *D. melanogaster* genome (Flybase R6.10) with Tophat version 2.1.0. Gene and read quantification were conducted using HTSeq version 2.7.6^60^, with reads under a quality score of 10 being discarded. Differential expression analysis was carried out using the limma package from R/Bioconductor^61^. Genes with counts per million (CPM) greater than 1 were retained. p values were adjusted using the Benjamini-Hochberg^62^. The overrepresentation analysis was performed by using ClusterProfiler package (version 4.12.6) in R, referencing KEGG, Reactome and Gene Ontology databases^63^. The heatmaps were generated from the scaled log2CPM values using tidyheatmaps R package.

### TaDa sequencing and analysis

Developmentally staged 1^st^ instar larvae were collected 24 hours after egg laying and maintained at a defined density on low sugar diet (LSD) plates at 25°C. Late second instar larvae expressing either *UAS-Dam-CtBP, UAS-Dam-Pros*, or *UAS-Dam* alone under the control of *Pros-Gal4*^*ts*^ driver were transferred to high sugar diet (HSD) plates for a 2-hour preadaptation at 25°C. Following this, larvae were moved to 29°C for 16 hours to induce transgene expression. A total of 75 larvae per sample were dissected in PBS and immediately snap-frozen in liquid nitrogen. Isolation and amplification of methylated genomic DNA from the larval guts was performed as described earlier^64^. Methylated DNA was further fragmented by sonication to achieve the size of 400 bp. Sequencing of methylated DNA, which passed all quality control checks, was done by Illumina.

The raw sequencing reads were aligned to the *Drosophila* genome (Dmel_r6.43) and analyzed using the pipeline from the Marshall lab (https://owenjm.github.io/damidseq_pipeline/) with some modifications. To minimize the effect of nonspecific methylation commonly found in accessible genomic areas, the binding profiles of DAM-fused proteins of interest were normalized to DAM-only DNA binding. The normalization was performed using the kernel density estimation of the log2 GATC ratio. Peaks that passed a stringent false discovery rate (FDR) of less than 0.01 were identified and mapped to the closest transcription start site. Peaks with at least a one-base pair overlap that were present in 4 out of 5 *Pros > UAS-Dam-CtBP* or 2 out of 3 *Pros > UAS-Dam-Pros* replicates were pooled together and identified as sites of true binding. Consequently, the width of the pooled peaks represented the total combined areas of the individual replicates. Finally, the consolidated peaks were analyzed across datasets to identify common target genes for CtBP and Prospero and significantly enriched pathways. For visualization, control-subtracted BigWig files were generated by using the bigwigCompare tool from the deepTools suite^65^ version 3.5.5. Subtracted replicate tracks were merged with the UCSC bigWigMerge to a bedGraph, averaged across replicates, and converted back to bigwig. Visualization of the resulting tracks and comparisons between conditions was carried out using the R package karyoploteR^66^ version 1.30.0. Custom CNET plots were generated using the igraph and ggplot2 R packages. Pathway gene annotations were based on KEGG, and direct targets were highlighted using data from targeted DamID experiments.

### *In silico* modelling of CtBP and Prospero interaction

The structure of Prospero in complex with dCtBP was predicted using the AlphaFold2 Multimer v3 model^31^, implemented via the ColabFold pipeline^67^. Due to the considerable length (1,703 residues) and largely disordered nature of Prospero, we employed a fragmentation-based approach specifically designed for identifying protein-peptide interactions^33^, enabling us to pinpoint the dCtBP-binding motif within Prospero. This strategy was further enhanced using the refined structural confidence metric, actifpTM^34^. ColabFold was executed locally with MMseqs2 used under default settings to generate multiple sequence alignments^68^. Structural inference was performed using 10 recycles, producing five models, which were subsequently subjected to relaxation to improve geometry and stereochemistry.

### Site-targeted mutagenesis of CtBP

To guide experimental validation of the dCtBP-Prospero interaction, alanine-scanning mutagenesis of dCtBP was designed using the Rosetta Flex ddG method^69^. The input structure consisted of the dCtBP substrate-binding domain bound to the Prospero-derived “ALSLV” motif. Key pocket residues of dCtBP (C38, M42, T50, V51, A52, F54, and H63) were individually mutated to alanine, while allowing backbone and side-chain flexibility in the bound motif chain. Flex ddG calculations were performed with 35,000 backrub trials, a maximum of 5,000 minimization iterations, and an absolute convergence threshold of 200. Final ΔΔG scores were reweighted using a generalized additive model (GAM), as described in the original Flex ddG protocol.

### Statistical analysis

Statistical analyses were performed in GraphPad Prism 10.2.0 and R/Bioconductor. For parametric data, two-tailed t-tests or one-way analysis of variance (ANOVA) followed by Tukey’s post hoc test were applied. For nonparametric data, the Kruskal-Wallis test or Wilcoxon rank-sum (Mann-Whitney U test) test was used, with multiple testing correction where applicable (FDR < 0.05). Pupariation and survival rate data were compared using the Log-Rank test. The specific statistical test used for each experiment is detailed in the corresponding figure legend.

## ACKNOWLEDGEMENTS

We thank Heini Lassila, Josef Gullmets, Nikola Grujic, Nisa Pitafi, and Helmi Grönlund for technical assistance. The Bloomington and Vienna *Drosophila* Stock Centers, Zurich ORFeome Project as well as Johannes Bischof are acknowledged for providing fly stocks and plasmids. We also thank Jan Veenstra and Mikio Furuse for sharing antibodies. This study was facilitated by the University of Helsinki *Drosophila* core facility (Hi-Fly), DNA Sequencing and Genomics Laboratory, and the Light microscopy unit (LMU) supported by Biocenter Finland and Helsinki Institute of Life Science.

