## Supplementary figures for "Metabolic control of enteroendocrine cell fate through a redox state sensor CtBP"

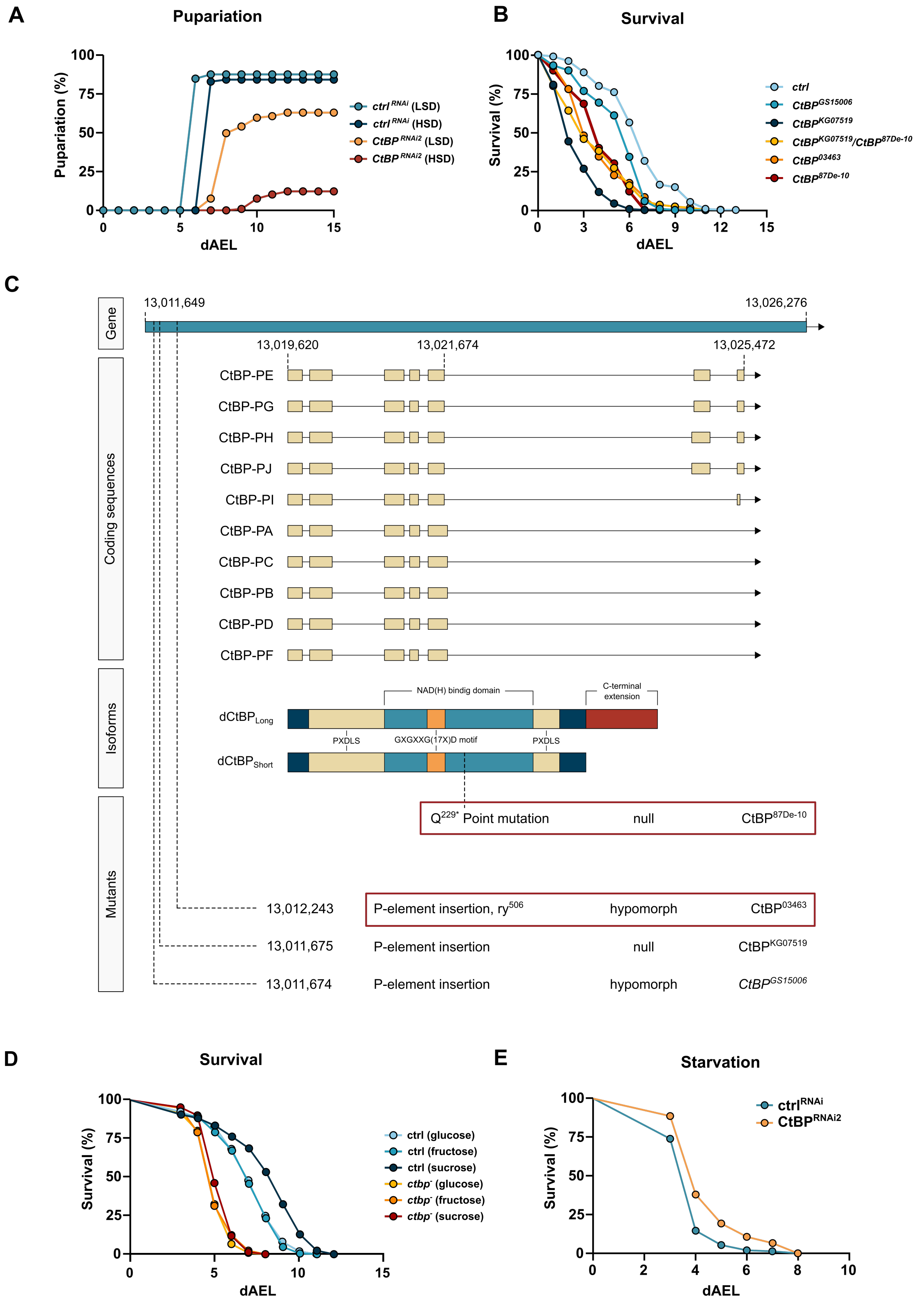


**Figure S1. CtBP regulates *Drosophila* energy metabolism and is necessary for sugar tolerance.**
**(A)** Whole-body knockdown of *CtBP* using an alternative RNAi line (*CtBP$^{RNAi2}$*) results in sugar intolerance. Pupariation rates for control and *CtBP$^{RNAi2}$* larvae on HSD were compared using the log-rank test ($\chi^2 = 71.9$, $P < 0.0001$). **(B)** Larvae carrying alternative *ctbp$^-$* mutant alleles (different from those presented in Figure 1D) lived significantly shorter on 5% sucrose (adjusted for multiple comparisons log-rank test, $\chi^2 = 262.9$, $P < 0.0001$). **(C)** Genetic characteristics of *ctbp$^-$* loss-of-function mutant alleles tested in this study. **(D)** *ctbp$^-$* mutants display comparable sensitivity to 15% glucose and fructose (log-rank test, $\chi^2 = 0.67$, $P = 0.41$). Their survival on a 15% sucrose is modestly but significantly different compared to the monosaccharides (adjusted for multiple comparisons, log-rank test, $\chi^2 = 10.48$, $P = 0.0048$). Overall, *ctbp$^-$* mutants exhibited significantly reduced survival compared to control animals (adjusted for multiple comparisons log-rank test, $\chi^2 = 584.3$, $P < 0.0001$) on a food with 15% dietary sugars. **(E)** Knockdown of *CtBP* using an alternative RNAi line (*CtBP$^{RNAi2}$*) increased starvation resistance of the larvae (log-rank test, $\chi^2 = 25.77$, $P < 0.0001$).

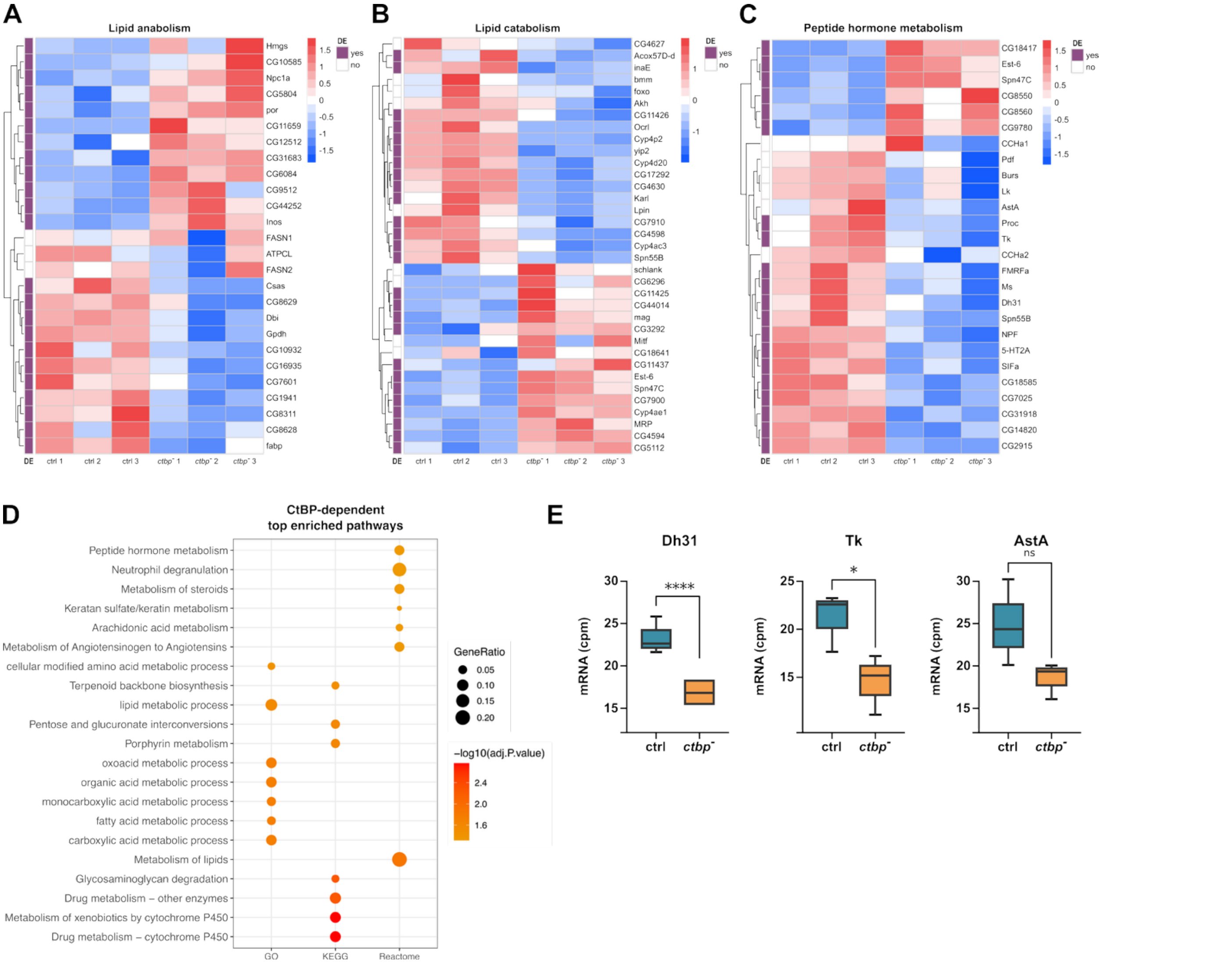


**Figure S2. CtBP acts in enteroendocrine (EE) cells to control organismal metabolism.**
*ctbp*-mutant larvae show dysregulation of genes involved in lipid anabolism **(A)** and catabolism **(B)** along with decreased expression and altered metabolism of peptide hormones **(C)**. The row-wise clustering was performed using Euclidean distance. Color key displays scaled log2 gene expression. (**D**) The top enriched pathways in *ctbp*-deficient larvae indicate a significant role for CtBP in peptide hormone and lipid metabolism. The color key represents the $\log_{10}$-adjusted P value, and circle size corresponds to the gene count in each pathway **(E)** Dh31, Tk, and AstA expression is impaired in *ctbp*-deficient larvae. *P = 0.0464;, **P = 0.011, $^{ns}$P = 0.129. Adjusted P-values were calculated using Benjamini-Hochberg correction from differential expression analysis.

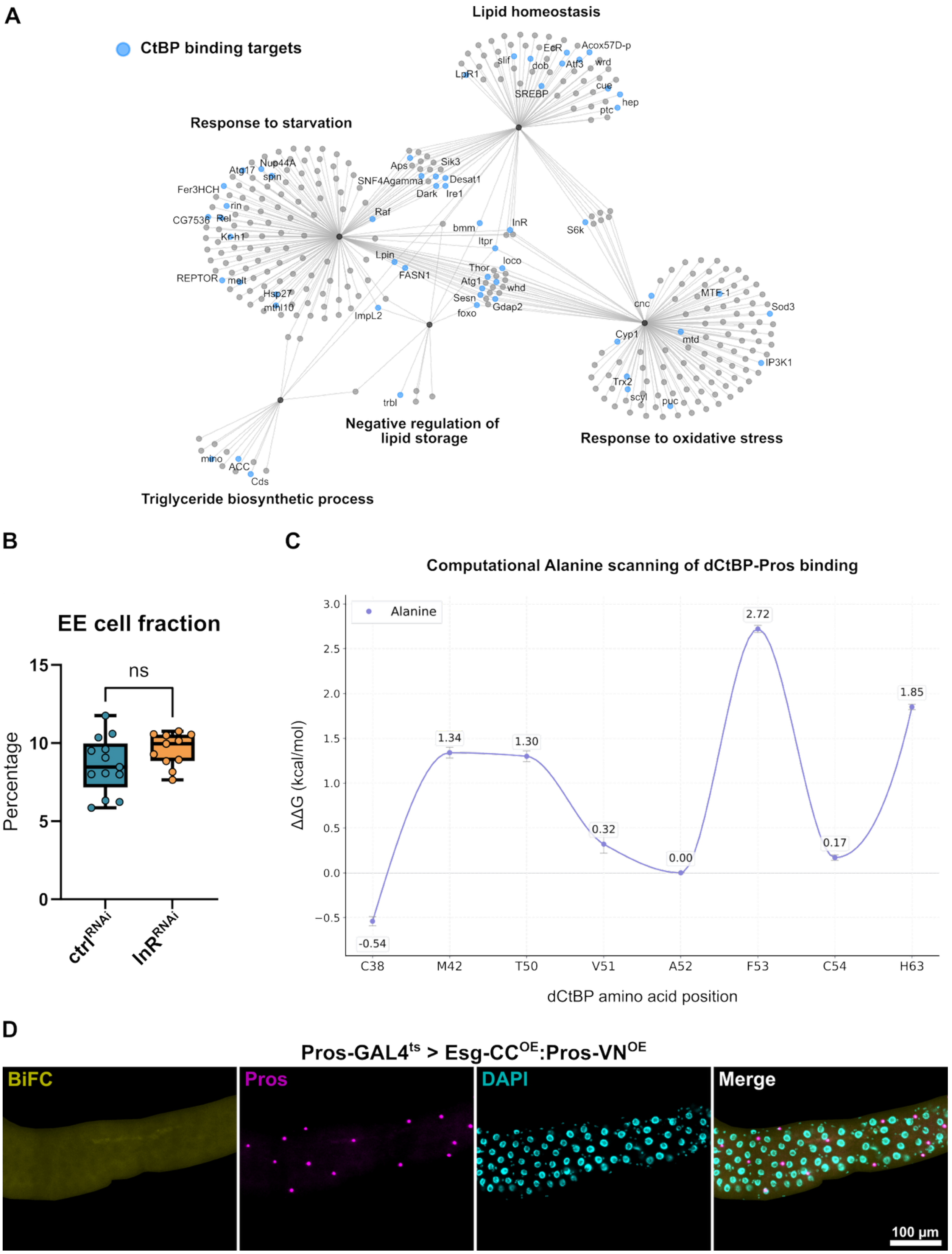


**Figure S3. CtBP controls EE cell fate through Pros-dependent and independent pathways.**
**(A)** CtBP genomic targets include regulators of lipid homeostasis and response to oxidative stress. CNET plot showing gene overlaps among selected pathways of CtBP targets identified by TaDa. Blue-colored nodes represent CtBP target genes within the pathway network. **(B)** Knockdown of *InR* in EE cells (Pros-GAL4>RNAi) does not affect the fraction of EE cells in larval intestine (unpaired t-test, ns indicates no significant difference in the studied parameter. **(C)** Computational scanning of dCtBP pocket residues for Pros binding reveals that the mutation of phenylalanine 53 to alanine (F53A) most significantly disrupts affinity with the Pros "ALSLV" motif ($\Delta\Delta G \approx 2.7$ kcal/mol). **(D)** Co-expression of Esg and CtBP BiFC fusion proteins in Prospero-driven EE cells does not produce fluorescence, serving as a negative control for the BiFC signal.
